# OmiGA: A Toolkit for Ultra-efficient Molecular Trait Analysis in Complex Populations

**DOI:** 10.1101/2024.12.19.629424

**Authors:** Jinyan Teng, Wenjing Zhang, Wentao Gong, Jiajian Chen, Yahui Gao, Lingzhao Fang, Zhe Zhang

**Author notes:** Corresponding Authors: **JT:** College of Animal Science, South China Agricultural University, Guangzhou, China, **LF:** Center for Quantitative Genetics and Genomics (QGG), Aarhus University, Aarhus, Denmark,;, **ZZ:** College of Animal Science, South China Agricultural University, Guangzhou, China. Equal contribution (Jinyan Teng, Wenjing Zhang).

## Abstract

Molecular quantitative trait loci (molQTL) mapping is one of the most popular approaches to systematically characterize functional impacts of genomic variants, leading to advanced understanding of the regulatory mechanisms underpinning complex traits and diseases. However, when applied to high-throughput molecular phenotypes, the existing molQTL mapping tools often implement simple linear models, overlooking complex inter-individual relatedness, leading to false positives and insufficient statistical power. Here, we introduce the **<u>Omi</u>**cs **<u>G</u>**enetic **<u>A</u>**nalysis toolkit (OmiGA), an ultra-efficient linear mixed model (LMM) based toolkit, for molQTL mapping in populations with complex relatedness. Both computational simulations and real data analyses demonstrated that OmiGA outperformed the existing popular tools regarding molQTL discovery power, fine mapping of causal variants, colocalization of molQTL and trait associations, and computational efficiency. In summary, we recommend OmiGA for molQTL mapping in populations with complex relatedness, for example, those in the Farm animal Genotype-Tissue Expression (FarmGTEx) project and family-based molQTL studies in humans.

## Introduction

The majority of trait-associated variants discovered by genome-wide association studies (GWAS) are non-coding and act through regulating intermediate molecular phenotypes (for example, gene expression and alternative splicing)^1,2^. Molecular quantitative trait loci (molQTL) mapping has been proposed to be a powerful approach to characterize the regulatory effects of genomic variants in natural populations, a step forwards to deciphering the regulatory circuitry of complex traits and diseases^3–7^. The conventional molQTL mapping often involves two steps: 1) is to perform a nominal association test and obtain the marginal effects of genetic variants on molecular phenotypes (e.g., gene expression) within the *cis*-region (e.g., 1 Mb around transcription start site, TSS) of the target gene; 2) is to conduct multiple testing correction to identify molGenes, i.e., genes with at least one significant *cis*-molQTL^7^. When analyzing these high-throughput molecular phenotypes (e.g., millions of gene expression across tissue and cell types), most existing molQTL mapping tools (e.g., fastQTL, tensorQTL, MatrixEQTL) implement simple linear regression models (LM) to reduce computational burdens^8–10^, resulting in the incomplete control of inter-individual relatedness, particularly in populations with complex relatedness (Fig. 1a-b).

**Fig. 1.**
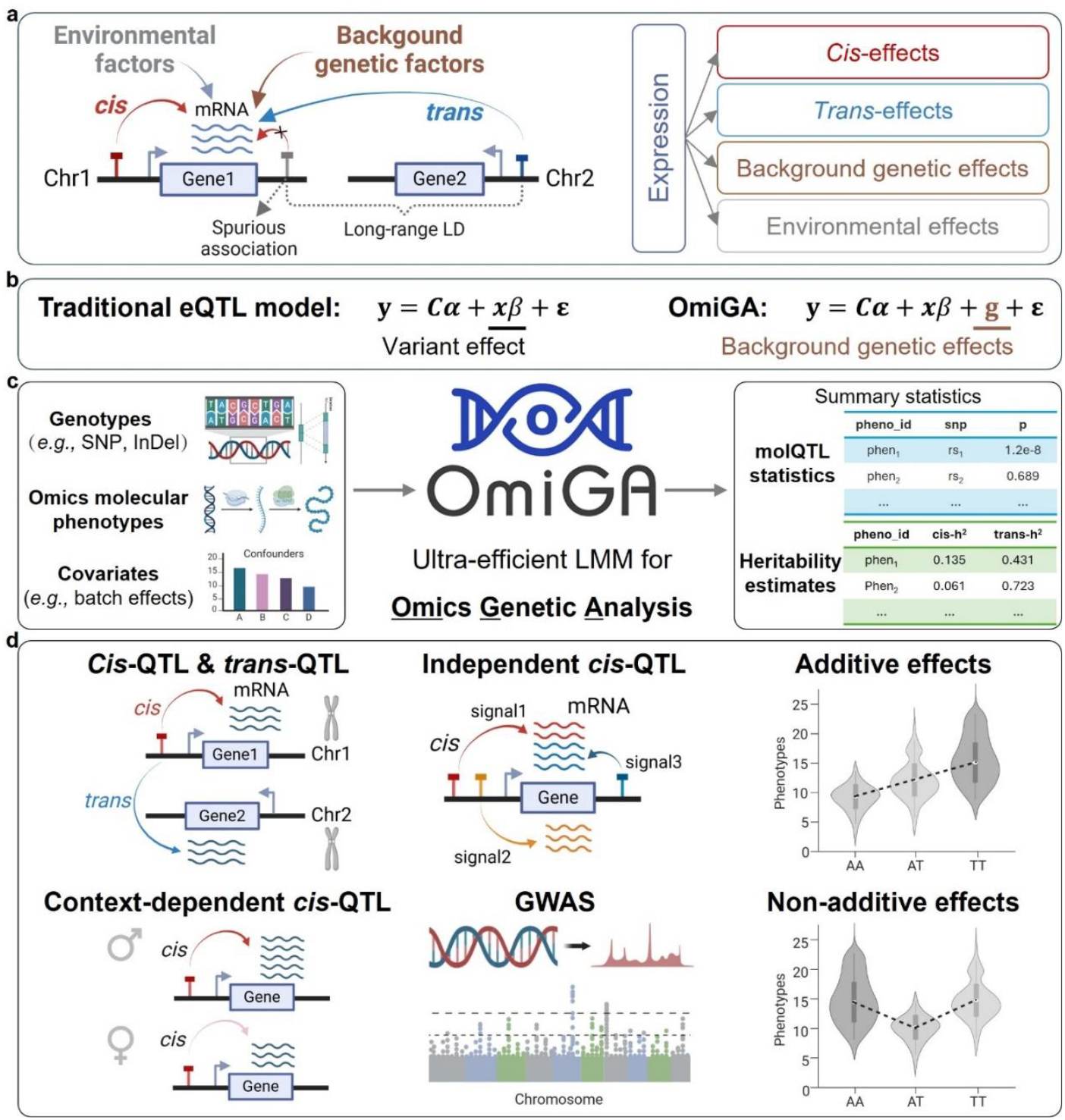
Overview of OmiGA for omics genetic analysis. **a**, The scenario of gene expression that is influenced by *cis*-variants, *trans*-variants, background genetic factors, and environmental factors. A suspicious non-causal *cis*-variant is shown downstream of Gene1, due to its linkage disequilibrium (LD) with the long-range *trans*-eQTL located on another chromosome. **b**, The statistical models of eQTL mapping for traditional tools and OmiGA. **y** represents the gene expression; ***C*** represents the covariates and *α* is its effects; ***x*** represents the genotype vector of tested variant and *β* is its effects; **g** is the component of background genetic effects; ε is the error term. The model of OmiGA incorporates a background genetic component to account for confounding genetic effects (such as long-range LD effects or relatedness) in eQTL mapping. **c**, The schematic diagram of input and output for OmiGA. OmiGA currently includes two key analysis modules: association study and heritability estimation. **d**, The main functions implemented in OmiGA include *cis*-QTL mapping, *trans*-QTL mapping, context-dependent *cis*-QTL mapping (i.e., interaction *cis*-QTL), conditional independent *cis*-QTL mapping, and genome-wide association study (GWAS) considering additive and/or non-additive effects. Additionally, OmiGA provides the estimation of *cis*-, *trans*-, and global heritability in both additive and/or non-additive effects.

Relatedness and population structure are crucial confounding variables that cannot be neglected^11,12^. To mitigate the impacts of relatedness among samples, a straightforward strategy is to analyze unrelated individuals only. This leads to a loss of statistical power of molQTL discovery, particularly in farm animals where most individuals are related due to the recent intensive breeding and limited effective population size. In the conventional GWAS, the linear mixed model (LMM) has been proposed as a standard approach to account for the complex relatedness among samples^13,14^. However, few molQTL mapping tools have implemented LMM due to their intensive computation required by estimating variance components for millions of molecular phenotypes via the restricted maximum likelihood (REML) algorithm^15,16^. Furthermore, existing molQTL mapping tools often ignore non-additive genetic regulatory effects, even though they are prevalent for molecular phenotypes^17,18^. Identifying variants with non-additive effects will allow us to better understand the underlying biology of complex traits^19^, particularly in farm animals^17^.

In this study, we developed an ultra-efficient and powerful **<u>Omi</u>**cs **<u>G</u>**enetic **<u>A</u>**nalysis toolkit (OmiGA) based on LMM for molecular QTL mapping. By conducting extensive simulations, we demonstrated that OmiGA outperformed all the existing molQTL tools regarding the statistical discovery power of molQTL and computational efficiency, especially for its robustness in controlling false positives in the presence of population stratification and complex relatedness. More specifically, OmiGA was up to ∼3.7 times faster than the commonly used LM-based tensorQTL (CPU version) in *cis*-eQTL mapping, and requires only ∼16% of the maximum memory usage compared to an LMM-based tool (APEX) for molQTL mapping, as well as is >26 and >13 times faster than the existing tools (GCTA^20^ and LDAK^21^) to estimate the global additive and dominant heritability of molecular traits, respectively. To showcase the utility of OmiGA, we mapped the *cis*-eQTL within four actual datasets sourced from pigs and humans. The pig datasets represent the population with complex relatedness, while human datasets represent the population consisting of unrelated individuals. We re-mapped *cis*-eQTL for 34 tissues in the pilot phase of the PigGTEx project, leading to a 64.54% increase in eGene discovery. Therefore, we recommend OmiGA (https://omiga.bio) for molQTL mapping in populations with complex relatedness, such as studies in the Farm animal Genotype-Tissue Expression (FarmGTEx) project.

## Results

### Overview of OmiGA

OmiGA is a toolkit designed for ultra-efficient molQTL mapping analysis, while accounting for population structure and relatedness, with technical details in Methods. The toolkit has two key modules (Fig. 1c-d): (1) LMM-based methods for molQTL mapping (e.g., *cis*-QTL, content-dependent *cis*-QTL, and *trans*-QTL) with additive and/or non-additive effects; and (2) additive and/or non-additive (*cis, trans*, and global) heritability partitioning for high-throughput molecular phenotypes. OmiGA requires three fundamental input data components: (1) complete or imputed genotype data; (2) a numerical matrix of molecular phenotypes with genomic annotations (e.g., TSS); and (3) covariates accounting for the batch effects and other hidden confounders (e.g., cell type composition in bulk tissue samples).

OmiGA performs molQTL analysis which comprises three steps: (1) The data rotation step, which involves constructing the genetic relationship matrix (GRM) using genotype data, performing the eigen decomposition of the GRM only once (with computational complexity of *O*(*n*^3^)), and rotating the genotype, phenotype, and covaraite using the eigen decomposition results with *O*((*m* +*p*+*c*)*n*^2^), where *n* represents the sample size, *p* is the number of molecular traits, *c* is the number of covariates, and *m* is the number of variants; (2) The nominal association test step, which involves estimating molQTL effects, 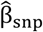 and 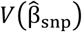), using a REML-free accelerated iterative dispersion that based on the newly developed AIDUL algorithm (see Methods)^22^. The computational complexities of AIDUL algorithm is lower, at *O*(*tpnc*^2^ + *mpc*^2^), compared to its original version (i.e., iterative dispersion update to fit LMM, IDUL^22^), which is *O*(*tmpnc*^2^), with the iteration number denoted as *t*. Additionally, it computes the association *P*-value for each variant across all molecular phenotypes using Wald test statistics 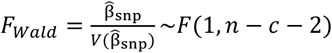; (3) The multiple-testing correction step, which involves two optional strategies that includes permutation-free aggregated Cauchy association test (ACAT)^23^ and *P*-value-free FDR control method, implemented in ClipperQTL^24^. Compared to thousands of permutations in tensorQTL, ClipperQTL requires fewer permutations to achieve comparable statistical power of molQTL discovery. The permutation test step of ClipperQTL involves shuffling phenotypes using two optional strategies to serve as response variables. This is followed by estimating partial correlation coefficients between the permuted phenotypes and genotypes using the proposed approximate partial correlation coefficients (APCC) method, which assumes that the relatedness contribution is negligible for the permuted phenotypes (as detailed in Methods). Ultimately, it reports significant molQTL associations under a given false discovery rate (FDR) threshold (i.e., 5% FDR).

To detect molQTL with dominant effects, we have implemented four optional statistical models (Methods). The computational procedures are similar to those used for additive molQTL mapping above, except when employing models that concurrently incorporate both additive and dominant GRMs (i.e., LMM(d + A + D)and LMM(a + d + A + D)). These models estimate additive genetic variance 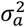, dominant genetic variance 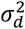 and residual variance 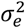 for each phenotype, necessitating iterative computation of the inverse of the variance-covariance matrix (**V**^−1^) tens of thousands of times (computational complexity of *O*(*tpn*^3^)). To address this computational challenge, we introduce the Pre-Allocated Eigen Decomposition (PAED) method to estimate the relevant variance components under the null model before conducting the nominal association test. This method reduces the computational complexity to *O*(κ*n*^3^ + *tpn*^2^), where κ is a finite value (i.e., 1000) dependent on the computational precision. PAED constructs a ‘prior’ combined GRM (cGRM) based on the variance component estimates and pre-computes the Eigen Decomposition, avoiding direct computing **V**^−1^ during the REML iteration. Subsequently, we optimize the statistical model by replacing the two GRMs with the cGRM, simplifying the model structure and enhancing computational efficiency. Ultimately, we leverage the AIDUL algorithm to efficiently calculate nominal association statistics.

We developed a specialized heritability estimation module for high-throughput molecular phenotypes within OmiGA. Unlike existing tools not optimized for molecular phenotypes, which repetitively estimate variance components for each phenotype, necessitating *O*(*tpn*^3^)computations for **V**^−1^ in every REML iteration, our approach is more efficient. We first perform Eigen Decomposition of the GRM once, obtaining **V**^−1^ through matrix-vector multiplication for each phenotype, resulting in a total computational complexity of *O*(*n*^3^ + *tpn*^2^)). For datasets with large sample size (e.g., >1000), we estimate variance components using two weighted least square regressions employing the IDUL algorithm. In the context of heritability models such as LMM(A_g_ + A_c_)and LMM(A_t_ + A_c_)with one GRM constructed from variants in the *cis*-region, we introduce a *cis* low-rank approximation (*cis*LRA) method. This reduces the **V**^−1^ computation complexity from *O*(*n*^3^)to *O*(*k*^3^)in the REML iteration, where *k* denotes the rank of the approximate low-rank matrix and *k* ≪ n. Furthermore, for heritability estimation involving both global additive and dominant GRMs (LMM(A_g_ + D_g_)), we leverage the PAED method to enhance computational speed and efficiency.

### Runtime and memory requirements

We evaluated the computational performance of OmiGA in comparison with existing tools that are specifically designed for molQTL analysis (i.e., LMM-based APEX and LM-based tensorQTL) in three datasets, including MAGE (Multi-ancestry Analysis of Gene Expression)^25^, GEUVADIS (Genetic European Variation in Disease)^26^ and GIAD042 (300 pigs)^27^. MAGE comprises 731 individuals from five distinct populations that encompasses a total of 19,539 expressed genes and 14,267,797 variants. GEUVADIS comprises 462 individuals from five distinct populations that encompasses a total of 17,703 expressed genes and 10,313,747 variants. GIAD042 comprises 300 individuals from three distinct populations that encompasses a total of 14,008 expressed genes and 15,495,927 variants (see Methods for details). We conducted comparisons of these tools at the same computing environment, which was allocated 120 GB of memory and 10 CPU threads (Intel Xeon CPU Max 9462) within a single server node. We then performed *cis*-eQTL mapping using OmiGA, APEX, and tensorQTL (CPU version). Since APEX does not support direct GRM calculation, we pre-computed the GRM using GCTA v1.94.1 and converted it to the sparse matrix format file that APEX accepts. Both OmiGA and tensorQTL utilized the same input files. We repeated five times to compare their average elapsed time and maximum memory usage.

We observed that OmiGA was up to ∼3.7 times faster than tensorQTL and required only slightly higher memory consumption (Table 1). To ensure that APEX could run properly, we set GRM elements to 0 for individuals with an estimated genetic relationship of more than 4^th^ degree (i.e., genetic relatedness < 0.044), as detailed in the previous study^28^. Although the runtime of OmiGA was 0.30 hours for GEUVADIS, which was slightly faster (∼1.8 times) than that of APEX based on a sparse GRM, OmiGA used much fewer computational resources than APEX (Table 1). More specifically, OmiGA required ∼14GB of memory to complete the whole analysis, only with approximately 16% of the maximum memory usage of APEX (∼90GB). Note that APEX cannot handle the MAGE and GIAD042 dataset with >14 million variants, despite the allocation of 500GB of memory.

**Table 1.** Comparison of runtime and memory usage of OmiGA, APEX, and tensorQTL.

| Dataset (sample size, variants) | OmiGA (LMM) | APEX (spLMM) | tensorQTL (LM) |
| --- | --- | --- | --- |
| MAGE (n=731, 14.3M) | 1.912 (28.516) | OOM | 2.293 (26.068) |
| GEUVADIS (n=462, 10.3M) | 0.210 (14.316) | 0.376 (89.900) | 0.775 (12.585) |
| GIAD042 (n=300, 15.5M) | 0.228 (14.697) | OOM | 0.799 (13.691) |

Elapsed runtime (hour) and maximum actual memory (GB) usage of OmiGA v1, APEX and CPU-version tensorQTL v1.0.9. The numbers in parentheses are memory usage. OMM means out of memory. All tests were performed in the same computing environment: 120 GB of memory and 10 CPU cores in one computer node.

Regarding heritability estimation, we compared OmiGA with two commonly used tools, namely GCTA v1.94.1 and LDAK v6. Because the GCTA and LDAK cannot automatically estimate heritability for tens of thousands of molecular phenotypes, we developed in-house pipelines. For global additive heritability estimation (i.e., Ag), we used the global additive GRM constructed by GCTA with *--make-grm* option as input for all three software. OmiGA showed >26 and >434 times faster than GCTA and LDAK, respectively (Table 2). For dominant heritability estimation (i.e., Ag+Dg), we used the dominant GRM constructed by GCTA with *--make-grm-d* option as input. OmiGA showed >13 and >204 times faster than GCTA and LDAK, respectively (Table 2). For *trans*- and *cis*-heritability estimation (i.e., At+Ac), OmiGA was faster than GCTA and LDAK, consistent with the global and dominant heritability estimation (Table 2). In addition, we observed that the estimates of variance components from OmiGA were nearly identical to those obtained from GCTA and LDAK, except for the estimation of dominant heritability using LDAK (Supplementary Fig. 1-3).

**Table 2.**
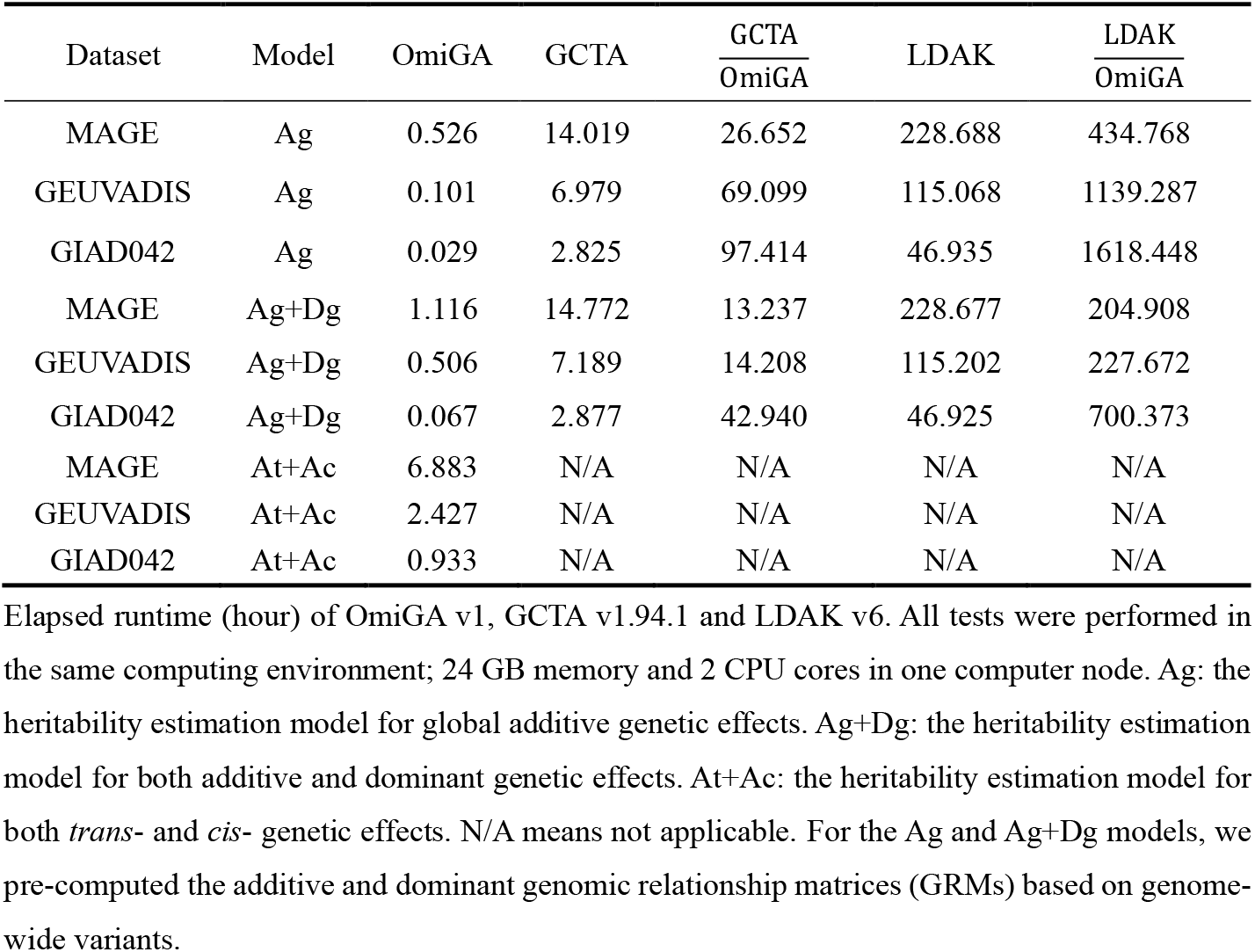
Comparison of runtime (hour) of OmiGA, GCTA and LDAK for heritability estimation of molecular phenotypes.

### OmiGA shows powerful performance in the simulations

Given the comparison results of resource requirements for different software above, we only considered OmiGA and tensorQTL, representing LMM and LM methods, respectively, to compare their statistical performances via extensive computational simulations. We then calculated the genomic inflation factor (λ), false discovery rate (FDR), F1-score, and true positive rate (TPR) (Methods) to evaluate their performances. One of the main aims of the simulations was to explore the influences of family relatedness on molQTL mapping. To mimic the population stratification and relatedness among individuals, we conducted simulations in two scenarios i.e., simulations with close relatedness individuals (SIM-CLOSE) and with unrelated individuals (SIM-UNREL) (see Methods). SIM-CLOSE used the GIAD042 dataset with close family relatedness, containing 300 pigs from three different populations (Fig. 2a). SIM-UNREL used the MAGE dataset containing 731 unrelated individuals from five different sub-populations (Fig. 2b). As expected, there were higher degrees of relatedness among individuals in GIAD042 than that in MAGE (Fig. 2a-b). Additionally, GIAD042 exhibited a much stronger inter-chromosome LD than MAGE (Fig. 2c-d, Extended Data Fig. 1, Supplementary Fig. 4-5).

**Fig. 2.**
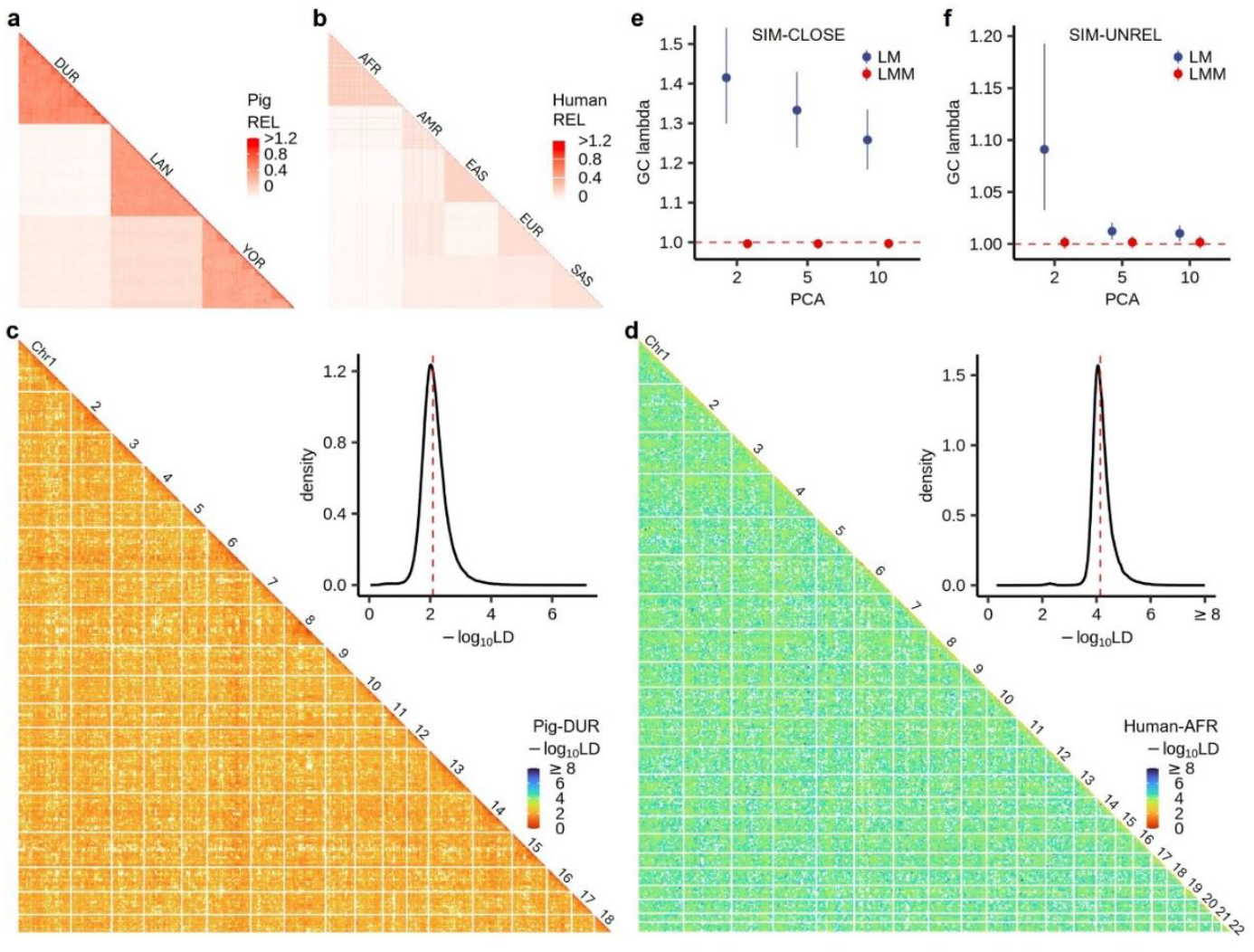
Population structure and inflation factor of test statistics on eQTL mapping in simulations. **a,b**, Heatmap of genomic relationship matrix (GRM) for all 300 pigs from GIAD042 (**a**) and all 731 individuals from MAGE (**b**) dataset. **c,d**, -log10-scaled linkage disequilibrium (LD, *r*^2^) and their distribution across the whole-genome variants for the Duroc (DUR) sub-population of GIAD042 (**c**) and Africa (AFR) sub-population of MAGE (**d**) datasets. We computed the whole genome-wide LD using GEAR (https://github.com/gc5k/gear2)^50^. The white lines in the heatmap splitted the different chromosomes. The red dotted lines in the density plot represent median values. **e,f**, Genomic control inflation factor (λ) for global eQTL mapping in the SIM-CLOSE (**e**) and SIM-UNREL (**f**) scenarios. SIM-CLOSE: simulated dataset with close relatedness individuals. SIM-UNREL: simulated dataset with unrelated individuals. The point and error bar represent the median value and quartile, respectively.

We simulated gene expression levels with causal variants randomly sampled from *cis*-regions (designated as eGene) or no causal variants (designated as non-eGene), under various genetic architectures. We quantified λ by performing genome-wide eQTL mapping, where λ was defined as the median chi-squared statistic divided by its expected value at the null simulations. We defined the FDR of *cis*-eQTL discovery as the proportion of non-eGenes with multiple-testing corrected *P* values less than a given threshold (e.g., 0.05). We defined the statistical discovery power of eQTL as the proportion of true positives (TPR) under a specified false discovery rate (FDR) (e.g., 5%). We mimicked the effect of population stratification by generating a mean phenotype difference between three pig populations and mimicked the effect of relatedness by specifying polygenic background effects at different levels (Methods).

We first compared the performance of OmiGA and tensorQTL to account for the population structure and relatedness. As expected, simulation results showed that the test statistics from LM were inflated across all scenarios with relatedness of different degrees (Fig. 2e-f, Extended Data Fig. 2). The genomic inflation factors in the test statistics of LM become smaller when increasing the number of genotype PCs in the model. The inflation in the test statistics for SIM-CLOSE population was stronger than that for SIM-UNREL population. This was due to the stronger long-range inter-chromosome LD in farm animals than that in humans (Fig. 2c,d). By contrast, there was almost no inflation in the test statistics of LMM with different numbers of genotype PCs across all simulations (Fig. 2e-f, Extended Data Fig. 2c-d), demonstrating the robustness of LMM-based OmiGA in accounting for long-range LD due to complex relatedness and small effective population size.

We further assessed the statistical power of OmiGA for *cis*-eQTL discovery, compared to tensorQTL. Based on the same number of genotype PCs, we quantified the FDR at a given significance level obtained from OmiGA and tensorQTL. We detected eGenes from OmiGA using ACAT pvalue < 0.05 at the gene level and those from tensorQTL using Storey’s qvalue < 0.05 which is used in the human GTEx^3,29^. In the SIM-CLOSE population, OmiGA and tensorQTL exhibited mean FDRs of 4.3% and 19.7%, respectively, and mean TPRs of 73.6% and 71.3% under 5% FDR (Fig. 3a). By contrast, in the SIM-UNREL population, OmiGA exhibited a slight improvement than tensorQTL, with the mean FDR of 3.7% and 6.0%, respectively (Fig. 3b). For genes exhibiting varying degrees of *cis*-heritability, OmiGA controlled the FDR more effectively and efficiently than tensorQTL regarding eGene discovery, particularly in the SIM-CLOSE population (Fig. 3c, Extended Data Fig. 3, Supplementary Fig. 7). The FDR of OmiGA could be controlled well using various multiple-testing approaches in both populations (Extended Data Fig. 4a-b). However, tensorQTL, using the approach of beta-approximate with Storey’s qvalue, could not control the FDR well in the SIM-CLOSE population (Extended Data Fig. 4c-d). Fitting more genotype PCs in tensorQTL reduces the FDR, which was in line with the observation regarding the genomic inflation factor. Additionally, the TPR of tensorQTL was consistently lower than that of OmiGA (Extended Data Fig. 5).

**Fig. 3.**
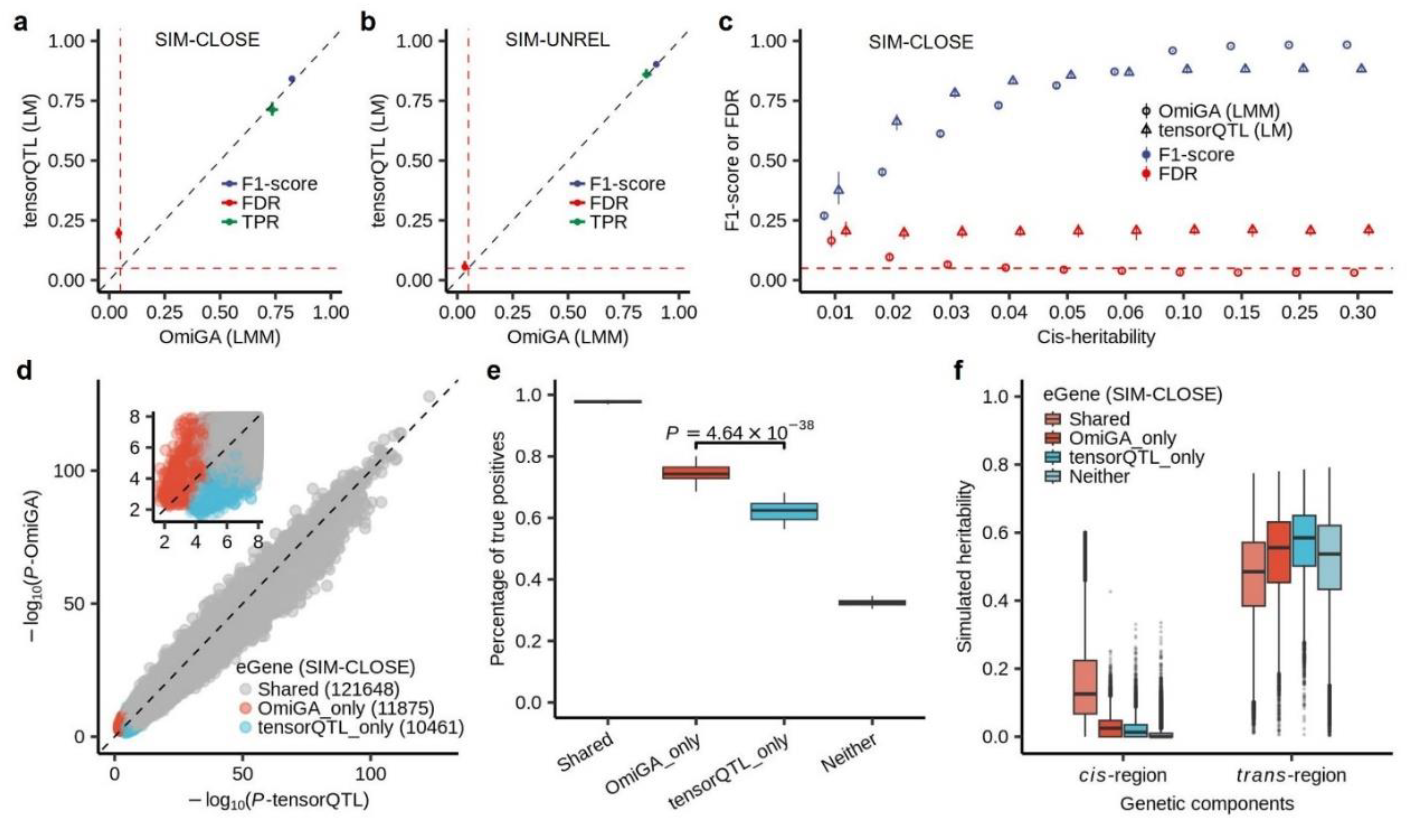
Performance of eQTL mapping using OmiGA and tensorQTL under different simulation scenarios. **a,b**, F1-score, false discovery rate (FDR), and true positives rate (TPR) of *cis*-eQTL mapping to identify eGene using linear mixed model (LMM) of OmiGA and linear model (LM) of tensorQTL in the SIM-CLOSE (**a**) and the SIM-UNREL (**b**). We defined eGene as the gene with at least one significant *cis*-eQTL. **c**, F1-score and FDR of eGene discovery for gene with different *cis*-heritability using LMM-based OmiGA and LM-based tensorQTL in the SIM-CLOSE. The points and the error bars represent the mean and the range, respectively in all panels. **d**, -log10-scaled *P*-values of lead *cis*-variants detected across 50 simulations. Shared: significant in both tools; OmiGA_only: significant in only OmiGA; tensorQTL_only: significant in only tensorQTL. **e**, Percentage of true positives of genes in different groups. Neither: no significant in either tool. The *P*-value was calculated from a two-sided Student’s *t*-test. **f**, True *cis*- and *trans* -heritability of genes in different groups.

To examine the characteristics of eGenes identified exclusively by OmiGA in the SIM-CLOSE population, we conducted a comparative analysis of the eGenes identified by both tools. To ensure a similar FDR between OmiGA and tensorQTL, we cc the results from tensorQTL under the Benjamini-Hochberg method and those from OmiGA based on ACAT pvalue < 0.05 at the gene level. The majority of eGenes were shared between OmiGA and tensorQTL (i.e., ‘shared’), evidenced by that mean TPR of shared eGenes identified by both tools was 97.8% across 50 simulations (Fig. 3d). However, there was still a certain proportion of true eGenes that were not discovered by both tools (Fig. 3e). The average percentage of true positives among eGenes detected by OmiGA only (i.e., ‘OmiGA_only’) was notably higher than those detected by tensorQTL only (i.e., ‘tensorQTL_only’) (Fig. 3e). Furthermore, the OmiGA_only eGenes exhibited a higher *cis*-heritability and lower *trans*-heritability compared to tensorQTL_only eGenes (Fig. 3f). These results indicate that OmiGA effectively controlled for long-range confounding effects originating from *trans*-region and identified a greater number of true positives compared to tensorQTL.

Additionally, we assessed the robustness of OmiGA in detecting context-dependent eQTL effects (also named interaction eQTL, ieQTL). We simulated *cis*-eQTL with interaction effects by incorporating an interaction term, as described in the Methods. We observed that OmiGA identified a higher number of true ieGenes, when maintaining the same FDR as those using tensorQTL in the SIM-CLOSE population (Supplementary Fig. 8-9), with mean TPRs of 69.0% and 64.7% for OmiGA and tensorQTL, respectively. Additionally, OmiGA demonstrated an advantage over tensorQTL in the discovery of ieQTLs with small interaction effects (explained ≤0.1 of genetic variance) (Supplementary Fig. 9a). In the SIM-UNREL, there were no obvious differences in association test statistics between OmiGA and tensorQTL (Supplementary Fig. 9b).

### Application of OmiGA to real data

We employed OmiGA to perform eQTL analysis in three tissues (duodenum, liver, and muscle) of 300 pigs from the GIAD042 population and lymphoblastoid cell line (LCL) of 731 humans from the MAGE population. As expected, there was inflation in the test statistics of tensorQTL, but not for OmiGA (Fig. 4a), in line with our observations from simulations (Fig. 2e-f). This indicates that OmiGA can effectively correct for the inflation caused by relatedness among individuals.

**Fig. 4.**
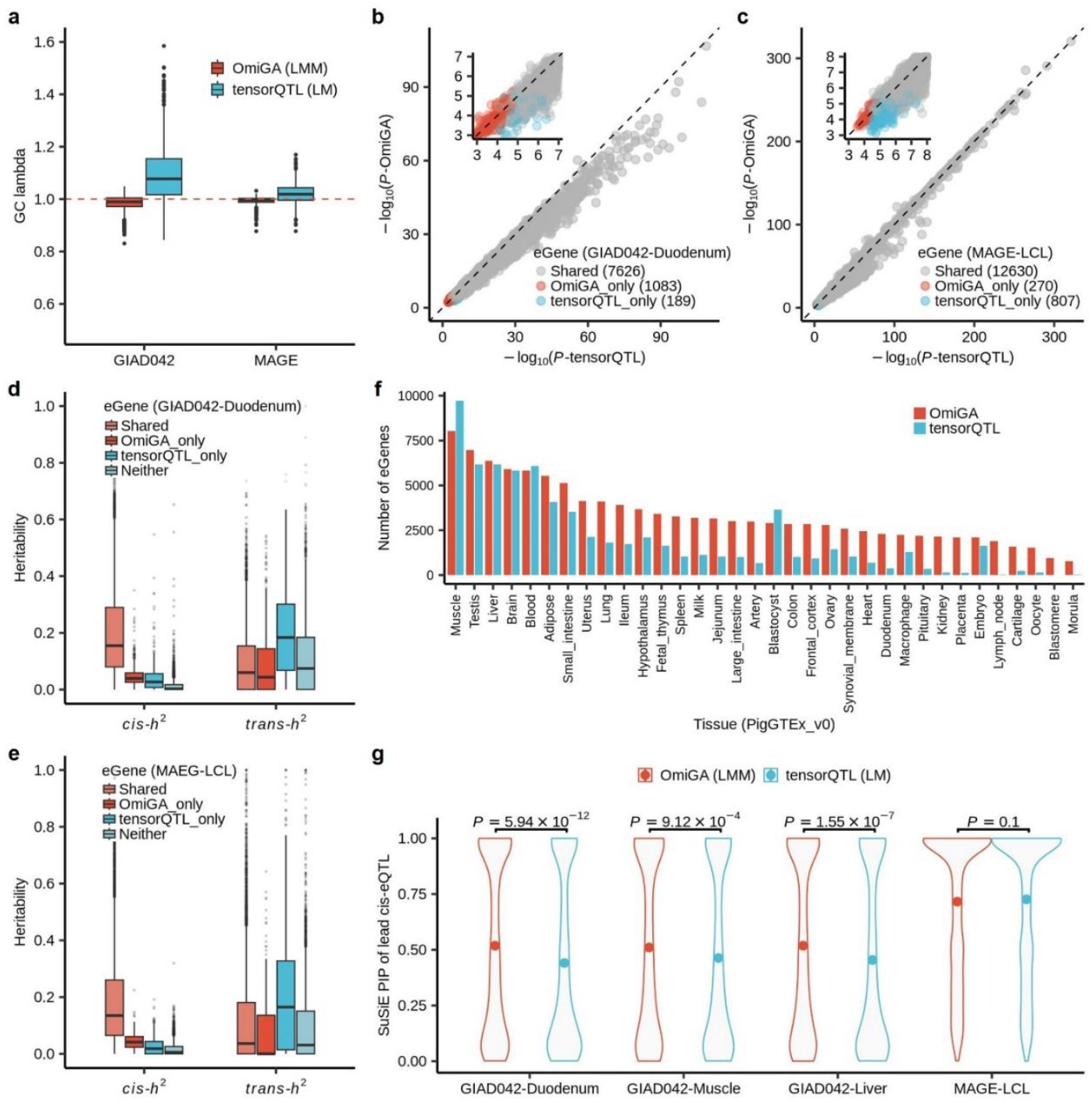
*Cis*-eQTL mapping using OmiGA and tensorQTL and fine-mapping in real datasets. **a**, Genomic control inflation factor of eQTL mapping. **b,c**, -log10-scaled *P*-values of lead *cis*-variants detected in duodenum from GIAD042 (**c**) and LCL from MAGE (**d**) dataset. **d,e**, Comparison of *cis*- and *trans*-heritability of eGenes discovered by OmiGA only and tensorQTL only in GIAD042 (**d**) and MAGE (**e**) dataset, respectively. **f**, Number of eGenes for 34 tissues from the pilot phase of PigGTEx project detected by OmiGA and tensorQTL, respectively. **g**, Comparison of causal posterior inclusion probability (PIP) for lead *cis*-eQTL obtained from OmiGA and tensorQTL in different tissues/datasets. PIP was computed using SuSiE-inf^51^. The point is the average of PIP. The *P*-value was calculated from a two-sided Student’s *t*-test.

We compared the discovery power of *cis*-eQTL between OmiGA and tensorQTL (Fig. 4b-c). Based on the experience of the simulation study above, to maintain a comparable FDR between OmiGA and tensorQTL for eGene identification, we employed the Benjamini-Hochberg method and qvalues with lambda=0.05 to detect eGenes in tensorQTL for GIAD042 and MAGE, respectively. Across the three tissues in GIAD042, OmiGA outperformed tensorQTL by identifying 1,083, 550, and 841 more eGenes detected in duodenum, muscle, and liver, respectively (Fig. 4b, Extended Data Fig. 6). Furthermore, OmiGA detected 270 additional eGenes in MAGE compared to tensorQTL (Fig. 4c). Consistent with the simulation findings, eGenes detected by OmiGA alone exhibited higher *cis*-heritability and lower *trans*-heritability compared to those detected by tensorQTL only (Fig. 4d-e, Extended Data Fig. 6), suggesting that OmiGA identified more true eGenes in these datasets. Moreover, OmiGA detected additional ieGenes in both datasets compared to tensorQTL. There are 512, 800, and 842 ieGenes that were only detected by OmiGA in the duodenum, muscle, and liver for GIAD042, respectively, and 133 ieGenes in LCL for MAGE (Supplementary Fig. 10). Building on these findings, we re-conducted *cis*-eQTL mapping for 34 tissues from the pilot phase of PigGTEx project^5^ using OmiGA. OmiGA achieved a 64.5% increase in eGene discovery compared to tensorQTL, with an average of 1,349 additional eGenes identified across 34 tissues (Fig. 4f). As anticipated, the pattern of *cis*- and *trans*-heritability of these eGenes discovered by OmiGA only was similar to those observed in simulations and the two real datasets above (Extended Data Fig. 6).

We employed SuSiE^30^ to perform fine mapping analysis for all eGenes. We derived the posterior inclusion probability (PIP) of the lead *cis*-eQTL of each eGene from OmiGA and tensorQTL. An average of 53% and 44% of tested eGenes from OmiGA and tensorQTL had credible sets of causal variants, respectively. Notably, the PIPs of lead *cis*-eQTL from OmiGA were significantly higher than those from tensorQTL (Fig. 4g), indicating that the *cis*-eQTL identified by OmiGA are more likely to be causal variants.

We then investigated whether the *cis*-eQTL discovered by OmiGA enhanced the statistical power for interpreting the GWAS associations of complex traits, compared to those identified by tensorQTL. We utilized GWAS summaries of 25 complex traits from PigBiobank to conduct colocalization analysis with *cis*-eQTL in duodenum, liver, and muscle^31^. The posterior probability of colocalization (PP.H4) using *cis*-eQTL from OmiGA was significantly higher than those from tensorQTL (Fig. 5a-b, Extended Data Fig. 7). For instance, *cis*-eQTL of the *TRIM52* gene in the duodenum identified by OmiGA colocalized (PP.H4=0.84) with the GWAS signal for lean meat percentage in pigs, whereas the PP.H4 for tensorQTL results was only 0.05 (i.e., no colocalization) (Fig. 5b). The colocalization signal around *rs324104102* in the GWAS was present in the OmiGA results but absent from the tensorQTL results. The Pearson’s correlation coefficient between the *P*-values of GWAS and *cis*-eQTL from OmiGA is 0.89, which was higher than that between GWAS and *cis*-eQTL from tensorQTL (Pearson’s *r*=0.79) (Fig. 5c-d). This discrepancy arises because the test statistics for variants downstream of *rs324104102* were inflation in the tensorQTL results, thereby reducing the correlation between the test statistics of GWAS and *cis*-eQTL.

**Fig. 5.**
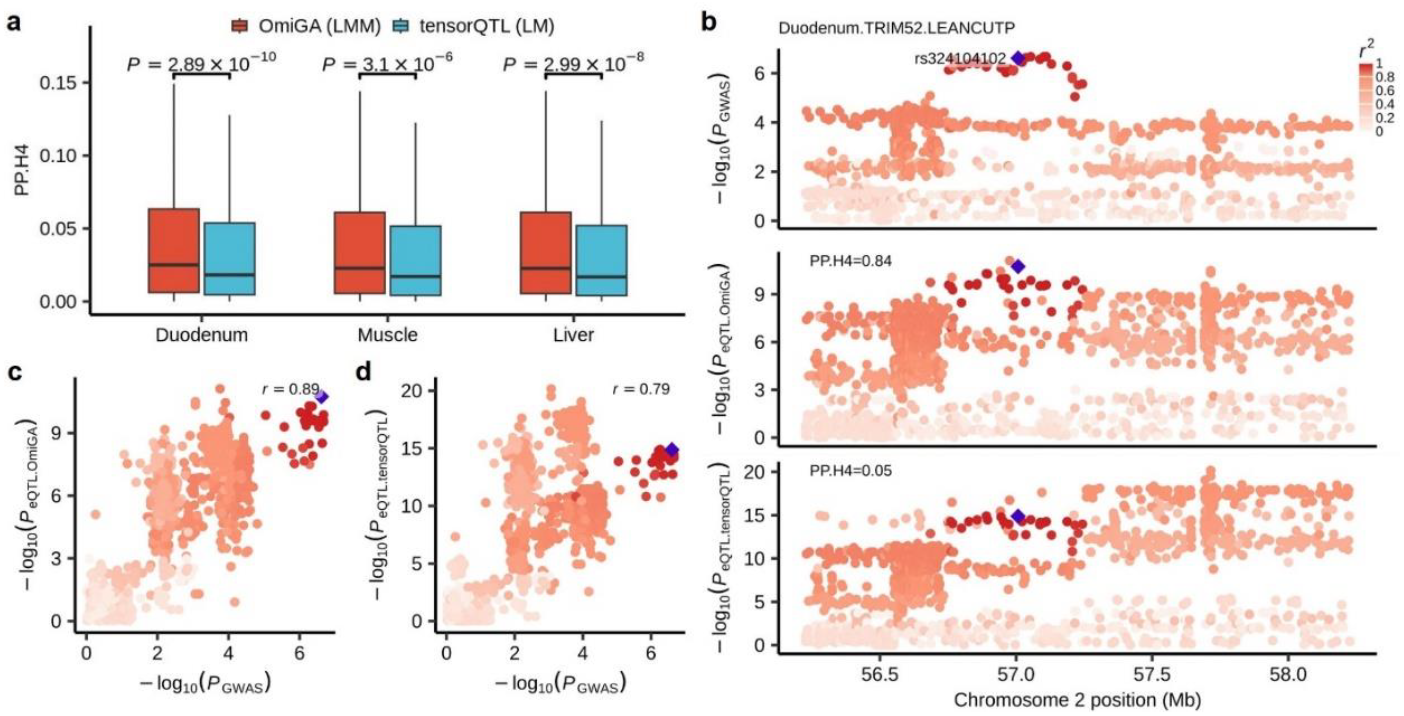
Colocalization of GWAS and *cis*-eQTL for GIAD042 dataset. **a**, Comparison of the posterior probability of colocalization (PP.H4) of GWAS and *cis*-eQTL that calculated from OmiGA and tensorQTL. We computed the PP.H4 using Coloc v5.2.3^52^. The *P*-values are obtained from a two-sided Student’s *t*-test. **b**, Colocalization of *cis*-eQTL of *TRIM52* in the duodenum and GWAS locus of lean meat percentage. **c,d**, Pearson’s correlation (*r*) between *P*-values of GWAS test statistics of lean meat percentage (LEANCUTP) and those of *cis*-eQTL from OmiGA (**c**) and tensorQTL (**d**).

We further utilized OmiGA to conduct *cis*-eQTL mapping in lymphoblastoid cell lines of all 731 individuals from the MAGE population. The performance of OmiGA was comparable to that of tensorQTL. This was because the MAGE cohort comprises unrelated individuals, rendering the use of LLM less necessary. Full summary statistics for LMM-based *cis*-eQTL of gene expression levels in the three tissues from the GIAD042 population, 34 tissues from the PigGTEx project, and LCLs from the MAGE population can be accessed publicly through the OmiGA portal at https://omiga.bio.

## Discussion

We have developed OmiGA, a powerful and efficient omics genetic analysis toolkit based on LMM that requires significantly lower system resources than existing tools. This makes it ideal for conducting molQTL mapping and heritability analyses on high-throughput molecular phenotypes within large-scale datasets, even when related individuals are present. In populations with relatedness, OmiGA exhibits superior robustness compared to LM-based tools. The efficiency and strength of OmiGA can be attributed to three key factors: (1) Optimization for resource-intensive omics high-throughput molecular phenotypes, reducing duplicate calculations and memory allocation; (2) Utilization of accelerated algorithms like AIDUL for fitting LMMs, which outperform existing algorithms in association study tools; and (3) Implementation of efficient methods for multiple testing correction, including the permutation-free ACAT method and the fast permutation Clipper method. Through simulations, OmiGA has demonstrated high robustness and statistical power in discovering *cis*-eQTL. Real data applications have shown that OmiGA identifies more reliable *cis*-eQTL, increasing the probability of being causal and colocalizing with complex traits. This suggests that using OmiGA for *cis*-eQTL mapping can facilitate GWAS fine-mapping. OmiGA efficiently handles datasets from both related and unrelated populations, providing precise association statistics for molQTL and facilitating downstream genetic analysis of complex traits.

Our research and previous studies have confirmed the effectiveness of LMM in addressing ancestry and population stratification, a fact substantiated by GWAS and molQTL studies^32–34^. However, the computational demands of LMM have limited its application in high-throughput molecular phenotype studies. For example, in a dataset with 20,000 genes, each having 10,000 *cis*-variants, the nominal association test in *cis*-QTL mapping would require a substantial number of variance components estimations and one hundred million association tests, leading to significant time and memory consumption. To expedite the estimation of variance components, existing association analysis tools use approximation methods that estimate variance components using REML algorithms under a null model excluding genetic variants^13,35^. Some methods attempt to simplify variance component estimations by adjusting covariates from phenotype values beforehand and employing the adjusted phenotype for association testing. However, this can reduce power, especially in multi-ancestry designs where the correlation between genetic effects and covariates varies. Hence, pre-adjusting phenotypes is not recommended in such scenarios. To overcome these challenges, we introduced a REML-free algorithm called AIDUL (see Methods) for fitting the LMM in molQTL mapping, requiring only two weighted least squares in each iteration. This markedly reduces the runtime compared to existing algorithms^22^. With this advanced algorithm optimization, OmiGA is suitable for conducting GWAS in large-scale populations as well.

The superior power of OmiGA over tensorQTL originates from the LMM is more adept handling for relatedness and population structure compared to the LM. This distinction is particularly pronounced in farm animal populations, where kinship relationships are prevalent, or human populations with significant degrees of relatedness^36^. While LM commonly employs PCs derived from genotype data to correct potential population structure, it falls short to capture the complex genetic relationships stemming from intricate family-based structures. Even when LM incorporates numerous genotype-derived PCs, as demonstrated in simulations, it inadequately adjusts for confounding relatedness (Fig. 2e-f, Extended Data Fig. 4c-e). In contrast, OmiGA, based on LMM, exhibits lower FDR and higher power. For robust LMM-based association analysis, it is recommended to include a few PCs as covariates in OmiGA^37,38^. Similar to GWAS for complex traits, in *cis*-eQTL mapping, it is recommended to assess the adequacy of adjusting for population structure and relatedness using the genomic inflation factor.

In molQTL mapping, where multiple-testing corrections are essential across variant-trait combinations, the correction method significantly impacts the power for molQTL discovery. While permutations are the gold standard for accurate multiple-testing correction, they come with a high computational cost. Various strategies have been developed to enhance the efficiency of multiple-testing correction, such as GPU-accelerated operations in tensorQTL^9^ and innovative approaches like ClipperQTL^24^ and eigenMT^39^. To address the computational burden of multiple-testing, OmiGA has implemented a permutation-free method: the aggregated Cauchy association test (ACAT)^23^. ACAT assesses the significance of molecular traits by aggregating *P*-values from each tested variant, with runtimes depending on the number of variants for a trait and remaining constant with increasing sample size. Simulation-based benchmarks have confirmed the efficiency and effectiveness of ACAT in determining the significance of molecular phenotypes compared to other methods.

In practical applications of OmiGA, several caveats need to be considered. Firstly, ensure missing genotypes or phenotypic values are imputed beforehand using appropriate tools. For instance, Beagle^40^ and HYFA^41^ are efficient tools for imputing missing genotypes and gene expressions, respectively. Secondly, exercise caution if variants with extremely low MAF and phenotypic outliers (such as low-expressed genes) have not been excluded, caution should be exercised during molQTL mapping to avoid false positive associations. In OmiGA, a useful option ‘--rm-pheno-threshold’ has been implemented to identify and eliminate phenotypes with abnormal distributions. Thirdly, OmiGA incorporates covariates to adjust for known batch and hidden confounding effects. Hidden confounders can often be inferred from phenotypic data using methods for hidden variable inference (e.g., PCA^42^, SVA^43^, PEER^44^, HCP^45^). As previously mentioned^42^, we recommend PCA due to its robustness. OmiGA offers an efficient option ‘--dprop-pc-covar’ to compute PCs from the input data. Fourthly, OmiGA employs a global GRM to correct for global population stratification but does not consider the local ancestry of each locus. While past studies have suggested that adjusting for local ancestry should slightly enhance the power of molQTL discovery, most results are consistent irrespective of whether adjustments are based on a local or global ancestry^46–48^. Fifthly, OmiGA conducts *trans*-eQTL mapping by default using all variants located outside of the chromosome of the target gene. Nonetheless, *trans*-eQTL mapping is highly prone to false positives due to mapping errors that result in short sequences being incorrectly mapped to homologous regions of the genome^49^. Therefore, meticulous filtering of variants is essential for *trans*-eQTL mapping. In the future, OmiGA aims to enhance its capabilities for genetic analysis of high-throughput molecular phenotypes, encompassing omics integration, genetics profiling, and association study.

## Acknowledgements

We thank all the researchers who have contributed to the publicly available data used in this research. We thank the members of the FarmGTEx Consortium for their assistance in testing and reporting software bugs, with special appreciation to Bingjin Lin, Mian Gong and Di Zhu. We are grateful to the National Supercomputing Center in Wuxi for doing the numerical calculations in this paper on its supercomputer system. Figure 1a,c,d were created with BioRender.com.

## Author Contributions Statement

J.T., Z.Z., and L.F. conceived and supervised the study. J.T. designed the experiment and developed the methods and software tools. W.Z. developed and performed the simulations with the assistance and guidance from Z.Z. and J.T. W.G. conducted data processing and performance comparison with the assistance of J.T. and W.Z. J.T. and W.Z. wrote the manuscript with the participation of J.C., Y.G., L.F., and Z.Z. All the authors reviewed and approved the final manuscript.

## Competing Interests Statement

The authors declare no competing interests.

## Extended Data Figures and legends

**Extended Data Fig. 1.**
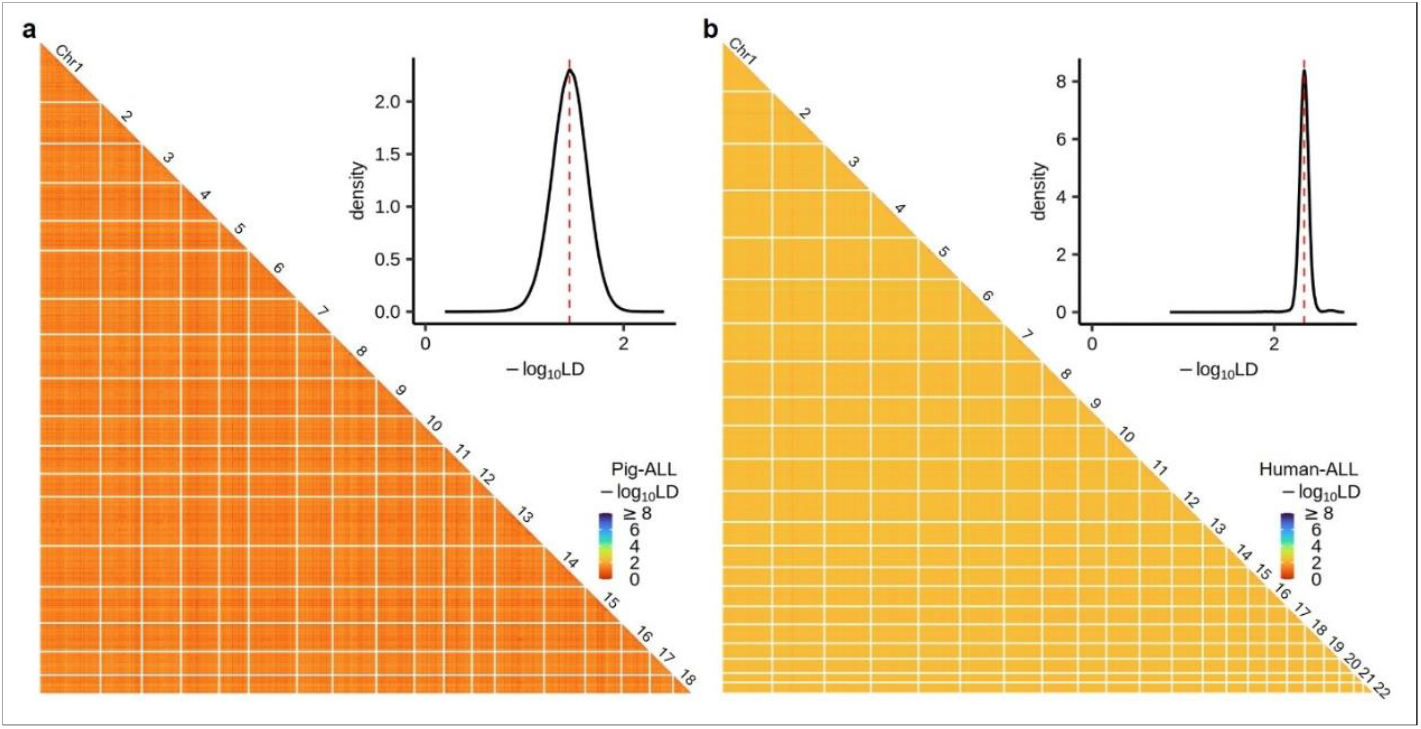
Linkage disequilibrium (LD) across all chromosomes. log10-scaled linkage disequilibrium (LD, *r*^2^) and their distribution across the whole-genome variants for GIAD042 population (**a**) and MAGE population (**b**). We estimated the global LD based on all individuals from each dataset using GEAR (https://github.com/gc5k/gear2)^50^. The white lines in the heatmap splitted the different chromosomes. The red dotted lines in the density plot represent median values.

**Extended Data Fig. 2.**
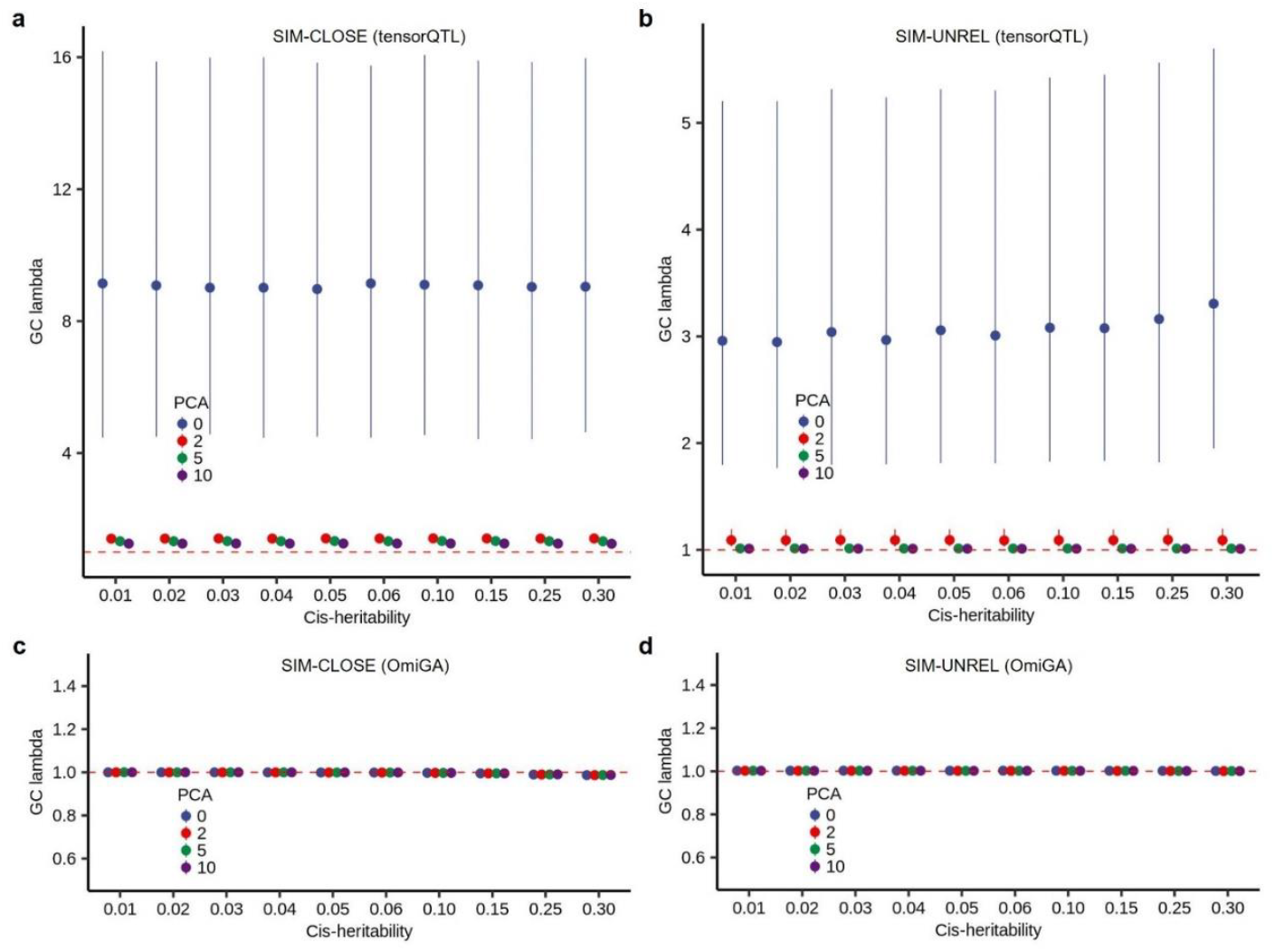
Genomic control inflation factor (GC lambda) of eQTL mapping for genes with different *cis*-heritability in different scenarios of simulations. **a,b**, eQTL mapping using a general linear model in tensorQTL for the SIM-CLOSE (**a**) and the SIM-UNREL (**b**) scenarios. **c,d**, eQTL mapping using a linear mixed model in OmiGA for the SIM-CLOSE (**c**) and the SIM-UNREL (**d**) scenarios. The point and error bar represent the median and quantile, respectively. The color of the point represents the number of genotype principal components (PCs) used in eQTL mapping model.

**Extended Data Fig. 3.**
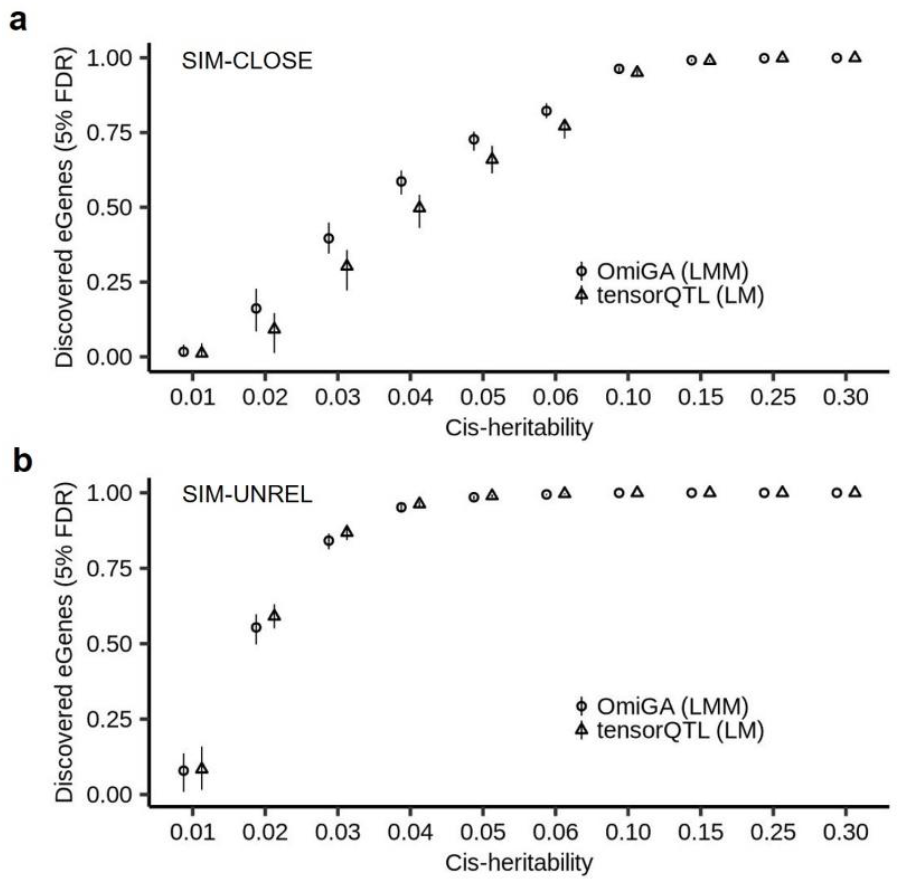
Proportion of simulated eGenes discovered by OmiGA and tensorQTL. For the *cis*-eQTL mapping results from the SIM-CLOSE (**a**) and the SIM-UNREL (**b**), we sorted the genes by the *P*-values at gene-level and summarized the proportion of eGenes discovered at 5% false discovery rate. The point and error bar represent the median and quantile, respectively.

**Extended Data Fig. 4.**
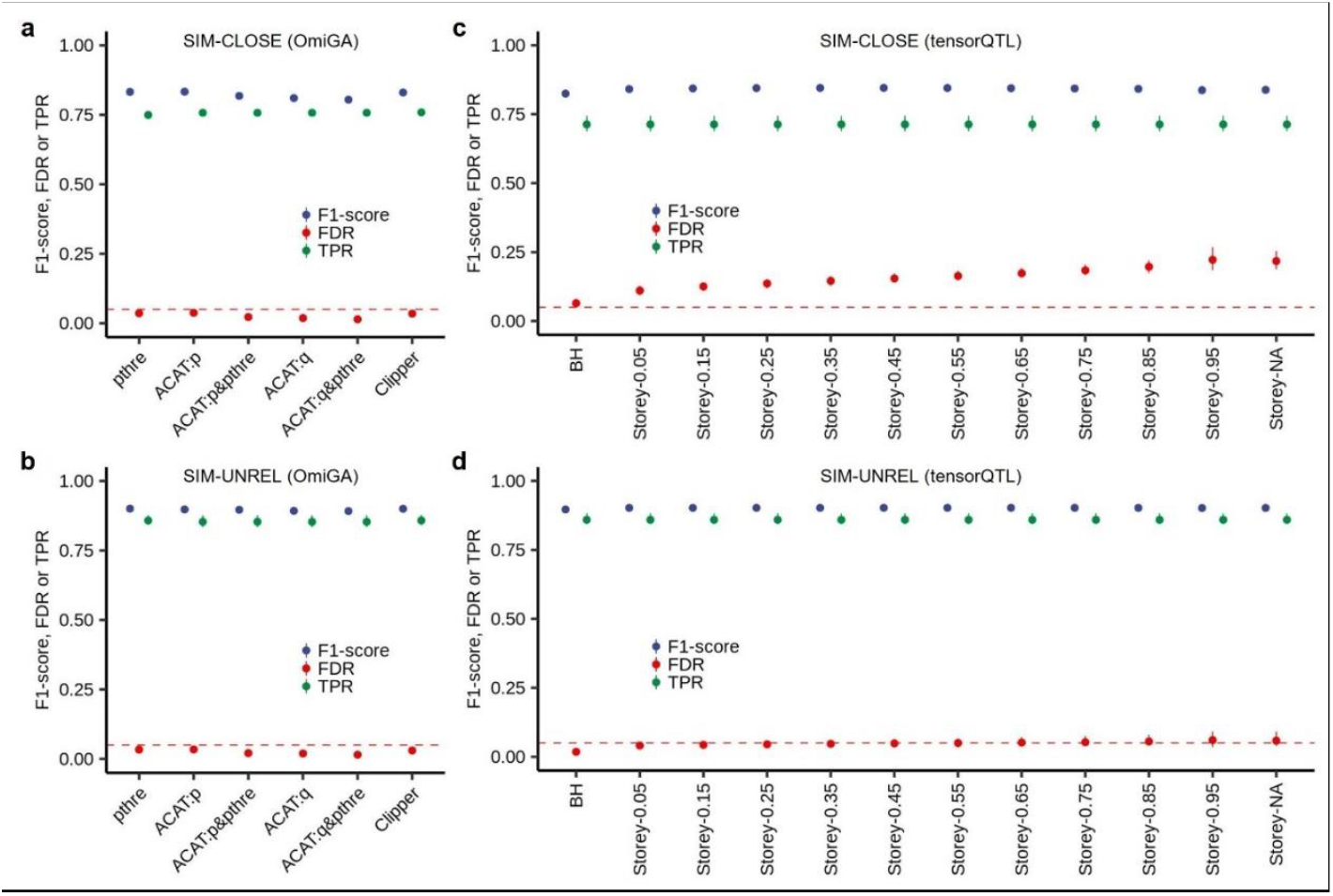
Comparison of multiple-testing approaches for eGene discovery. **a,b**, *Cis*-eQTL mapping using OmiGA in the SIM-CLOSE (**a**) and the SIM-UNREL (**b**). pthre: the nominal *P*-values of lead variants below the variant-level threshold obtained from permutations. ACAT: aggregated Cauchy association test. ACAT:p: the gene-level *P*-values obtained from ACAT < 0.05. ACAT:p&pthre: the combinations of ACAT:p and pthre. ACAT:q: the Benjamini & Hochberg (BH) adjusted gene-level *P*-values obtained from ACAT < 0.05. ACAT:q&pthre: the combinations of ACAT:q and pthre. Clipper: the gene-level *P*-values obtained from ClipperQTL < 0.05. **c,d**, *Cis*-eQTL mapping using tensorQTL in the SIM-CLOSE (**c**) and SIM-UNREL (**d**). BH: the BH adjusted gene-level *P*-values < 0.05. Storey-*: the gene-level Storey’s qvalues < 0.05 with different lambda parameters. The points and the error bars represent the mean and the range, respectively, in all panels.

**Extended Data Fig. 5.**
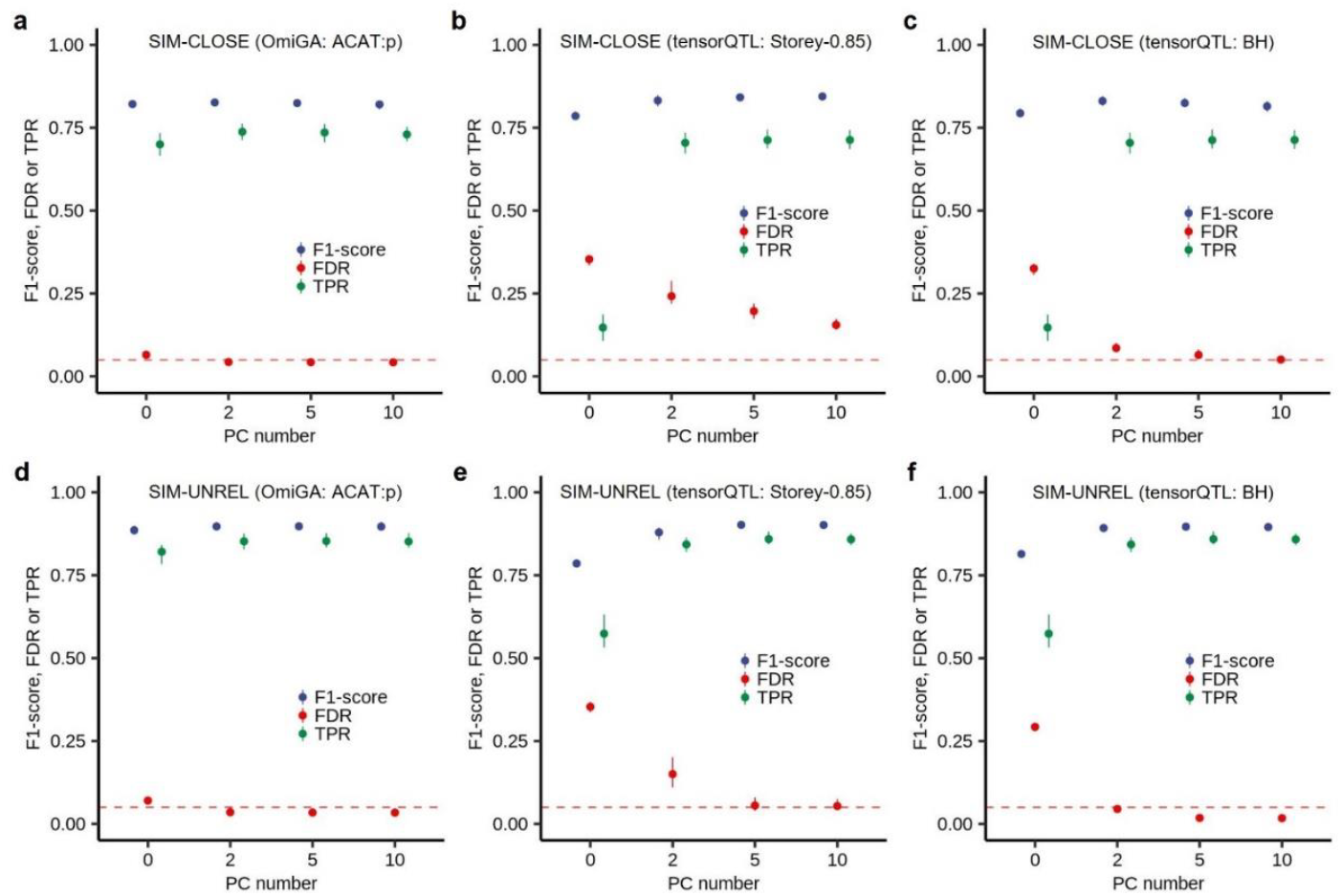
Comparison of different number of genotype principal component (PC) used for *cis*-eQTL mapping. **a-c**, *Cis*-eQTL mapping using OmiGA with ACAT:p (**a**) and tensorQTL with Storey-0.85 (**b**) and BH (**c**) in the SIM-CLOSE. Storey-0.85 is the same as did in GTEx v8. **d-f**, *Cis*-eQTL mapping using OmiGA with ACAT:p (**d**) and tensorQTL with Storey-0.85 (**e**) and BH (**f**) in the SIM-UNREL. ACAT: aggregated Cauchy association test. ACAT:p: the gene-level *P*-values obtained from ACAT < 0.05. Storey-0.85: the gene-level Storey’s qvalues < 0.05 with lambda=0.85. BH: the BH adjusted gene-level *P*-values < 0.05. The points and the error bars represent the mean and the range, respectively, in all panels.

**Extended Data Fig. 6.**
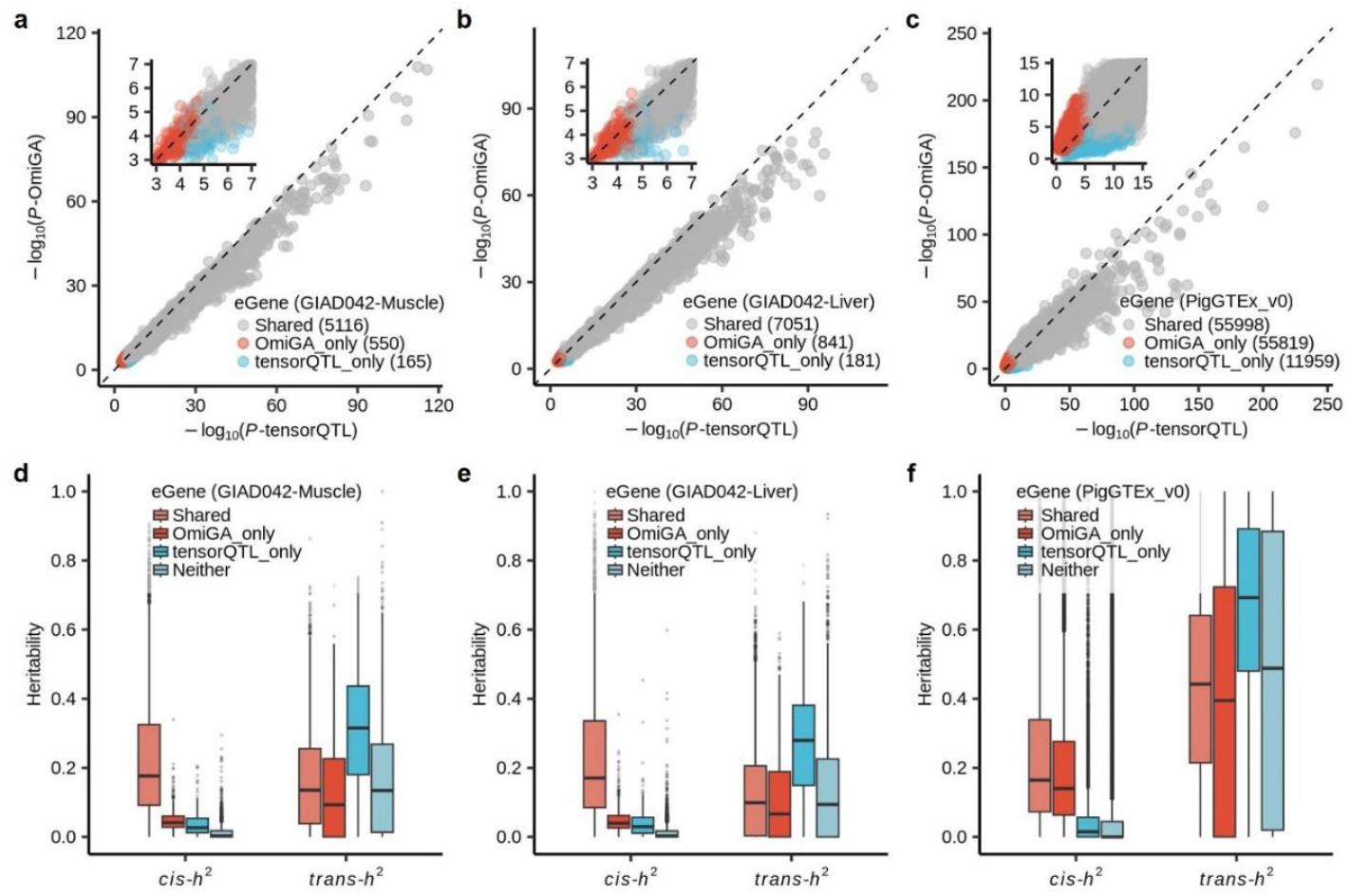
Comparison of eGenes discovered by OmiGA and tensorQTL. **a-c**, -log10-scaled *P*-values of lead *cis*-variants detected in muscle (**a**) and liver (**b**) from GIAD042 and 34 tissues (**c**) from the pilot phase of PigGTEx project. **d-f**, Comparison of *cis*- and *trans*-heritability of eGenes discovered by OmiGA only and tensorQTL only in muscle (**d**) and liver (**e**) from GIAD042 and 34 tissues (**f**) from the pilot phase of PigGTEx project, respectively.

**Extended Data Fig. 7.**
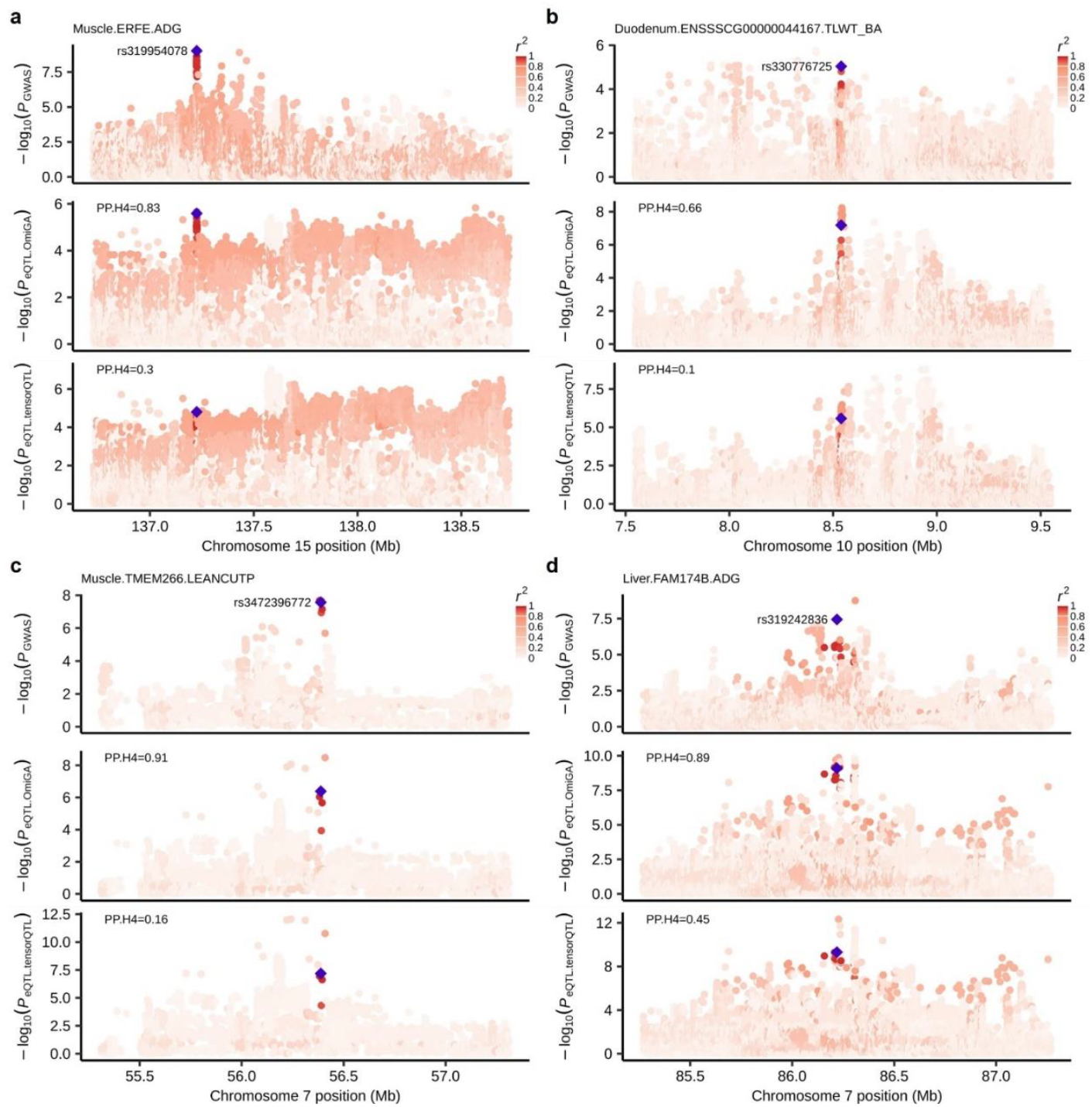
Examples of colocalization of GWAS and *cis*-eQTL for GIAD042 dataset. **a**, Colocalization of *cis*-eQTL of *ERFE* in muscle and GWAS locus of average daily gain (ADG). **b**, Colocalization of *cis*-eQTL of *ENSSSCG00000044167* in duodenum and GWAS locus of total litter weight of piglets born alive (TLWT_BA). **c**, Colocalization of *cis*-eQTL of *TMEM266* in muscle and GWAS locus of lean meat percentage (LEANCUTP). **d**, Colocalization of *cis*-eQTL of *FAM174B* in liver and GWAS locus of average daily gain (ADG). We computed the posterior probability of colocalization (PP.H4) using Coloc v5.2.3^52^.

## Online Methods

### Ethics

Not applicable because no biological samples were collected, and no animal handling was performed for this study.

### Linear mixed model for standard molecular QTL mapping

To detect molQTL with additive effects, we consider the following standard linear mixed model:

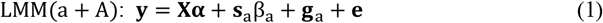

where **y** is an *n*×1 vector of standardized molecular phenotypes, **X** is an *n*×*c* matrix of customized covariates including a column of 1, **α** is the fixed effect corresponding to covariates, **s**_a_ is an *n*×1 vector of mean centered genotype values in additive coding (0/1/2 for AA/Aa/aa) for the genetic variant being tested, β_a_ is the variant’s additive genetic effect, **g**_a_ is an *n*×1 vector of total additive genetic effects with 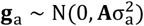 where 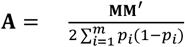 is an additive genetic relationship matrix (GRM), of which **M** is a matrix of mean centered genotypes for genome-wide *m* genetic variants and *p*_*i*_ is the MAF at *i*^th^ variant, **e** is *n*×1 vector of residuals with 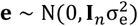 where **I**_*n*_ is an *n*×*n* identity matrix. To obtain association statistical magnitude, we employed exact and accelerated methods described below.

### AIDUL for association test

To accelerate the association test process, we propose an accelerated iterative dispersion update to fit linear mixed model (AIDUL) by modifying the IDUL (iterative dispersion update to fit linear mixed model) algorithm^22^. The computational complexities for these two algorithms are *O*(*tpnc*^2^ + *mpc*^2^)and *O*(*tmpnc*^2^), respectively, with the iteration number denoted as *t*. We first randomly select a small subset of variants for target gene and perform the IDUL process as below. The model (1) assumes **g**_a_ ∼ MVN_*n*_(0, *ητ*^−1^**A**), **e** ∼ MVN_*n*_(0, *τ*^−1^**I**_*n*_), where *τ*^−1^ is the variance of the residual errors 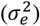, *η* is the ratio between the additive genetic variance 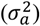 and 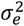, MVN denotes multivariate normal distribution. The eigen decomposition of **A** can be written as **QTQ**^′^ with **QQ**^′^ = **I**_*n*_, where *j*^th^ column of **Q** is an eigenvector, whose corresponding eigenvalue is the *j*^th^ diagonal element of the diagonal matrix **T**. Let **W** = (**X, s**_a_)and **b** = (**α**, β_a_), we can write the formula (1) as follows:

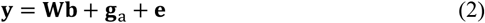

We rotate the formula (2) by multiplying **Q**^′^ to get

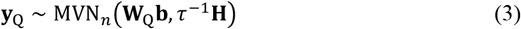

**y**_Q_ ∼ MVN_*n*_(**W**_Q_b, *τ*^−1^**H**)(3) where **y**_Q_ = **Q**^′^**y**, W_Q_ = **Q**^′^**W**, and **H** = *η***T** + **I**_*n*_ . We perform IDUL until *η* converges by repeating:

Step 0: assign a constant to *η*_*i*_ (e.g., 0.1) for the first iteration (*i* = 1) of IDUL.

Step 1: fit formula (3) using weighted least squares with weight **H**^−1^ to obtain the residual **r**.

Step 2: fit **r**^2^ ∼ MVN_*n*_(*µ* + diag(T)γ, *τ*^−1^**H**^2^)using weighted least squares with weight **H**^−2^ to obtain estimates 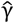 and 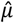.

Step 3: update 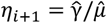 and repeat step 1 to 3 until 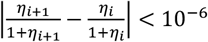 or the number of iterations achieve to the maximum iterations (e.g., 100).

After the *η* converges, we perform AIDUL process by obtaining the average estimates of *η*_*avg*_ for the subset of variants. We then let *η* = *η*_*avg*_ in formula (3) and obtain the effect estimates 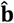 and its variance 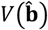, where 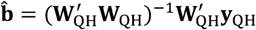 of which **W**_QH_ = W_Q_**H**^−1^ and **y**_QH_ = **y**_Q_**H**^−1^, 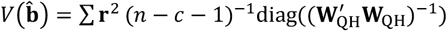. Under the null hypothesis, the Wald test statistics *F*_*Wald*_∼*F*(1, n −*c* − 1)and the *P* value of effect (β_a_) of tested genetic variant can be calculated.

### Multiple-testing corrections for *cis*-molQTL mapping

To adjust the multiple testing across molecular phenotypes in *cis*-molQTL mapping, we implement two approaches including ACAT^23^ and Clipper^24^.

ACAT is a permutation-free and computationally efficient *P*-value combination method. We compute the ACAT test statistic using a linear combination of transformed *P*-values by 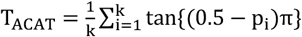, where k is the total number of variants for target gene, p_i_ is the nominal *P*-value of *i*^th^ variants tested in association test, and tan{(0.5 − p_i_)π is Cauchy distributed if p_i_is from the null distribution. Therefore, we then used T_ACAT_ to calculate the *P*-value of each molecular trait using the Cauchy cumulative distribution function to obtain the trait-level *P*-value.

Clipper is a *P*-value-free FDR control method to identify which phenotypes have significant *cis*-QTL. Clipper requires both measurements under the experimental and background conditions to determine the phenotype-level FDR. We define the phenotype-level partial correlation between phenotypes and lead variants observed in nominal association test as experimental measurements, and those from permutation test as background measurements for Clipper, where the lead variant is the most significant/correlated variant for a given phenotype or permuted phenotype.

To obtain the experimental measurements, we compute the partial correlation (*r*_*p*_) between **y** and **s**_a_ for lead variant using 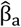 and 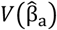 estimated from nominal association test (i.e., LMM(a + A)model (1)). The lead variant with the largest absolute partial correlation 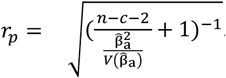. To obtain the background measurements, we implemented two approaches for permutation test including direct permutation and residual permutation schemes.

#### Direct scheme

We perform direct permutation test using the following model:

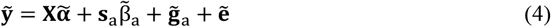

where 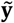 is an *n*×1 vector of molecular phenotypes being shuffling, 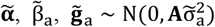, and 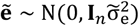 are fixed effect of covariates, genetic effect of tested variant, total additive genetic effects, and residuals, respectively, corresponding to permuted phenotype values. To reduce the computationally intensive in the permutation process, we assign zero to 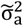 for permuted phenotypes assuming the genetic contribution is negligible. Therefore, the term 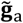 can be neglected and the approximate model for direct permutation is a simple linear regression model:

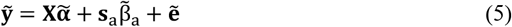

To eliminate the effects of covariates, we calculate the residualised phenotypes by 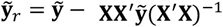, and the residualised genotypes by **s**_*r*_ = s_a_ − **XX**^′^**s**_a_(**X**^′^**X**)^−1^ . To efficiently compute the partial correlation, we standardize the 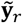 and **s**_*r*_ to have zero mean and unit sum of squares. Therefore, we can simplifies the calculation of the partial correlation as 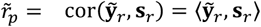 where 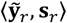 is the inner product between 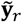 and **s**_*r*_.

#### Residual scheme

We perform residual permutation using ClipperQTL method^24^. Different from direct permutation scheme, residual permutation scheme first residualise phenotypes before shuffling step, and calculate 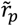 using 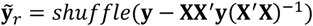as above. The residual scheme requires for data sets with large sample sizes (> 450 is recommended) and the direct scheme is suitable for a wide range of sample sizes.

After finishing permutation process, we can obtain a matrix 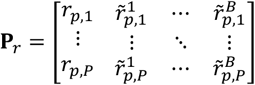 consisting of the maximum partial correlation coefficients for corresponding conditions, where the first column is experimental measurements, the second and subsequent columns are background measurements, subscript *P* is the number of tested phenotypes, and superscripts *B* is the number of permutations. As recommend by ^24^, *B* is set at 1000 by default for the direct scheme and 20∼100 for the residual scheme.

### Variant-level significance threshold

To identify the significance of variants for a target molecular trait, we first conduct a predetermined number of permutations by randomly shuffling the phenotype values across individuals. Subsequently, we derive the *P*-values for the lead variants from each of these permutations. The variant-level threshold for each gene is calculated as the 5^th^ percentile of permuted *P*-values of lead variants.

### LMM for interaction molecular QTL mapping

Extending the standard LMM(a + A)model (1), we test interaction effects between target genetic variant and interaction factors (e.g., cell-types or ancestry component) on molecular phenotypes using the following model:

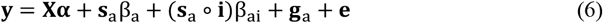

where ∘ is Hadamard product, **i** is an *n*×1 vector of the standardized interaction factor, (**s**_a_ ∘ **i**)is the interaction term between genotype and interaction factor, β_ai_ is the interaction effect of the genetic variant being tested, the rest of the terms the same as in LMM(a + A)model (1). Let **W** = (**X, s**_a_, **s**_a_ ∘ **i**)and **b** = (**α**, β_a_, β_ai_), based on IDUL method, we can obtain estimates 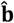 and 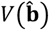 as in formula (3) as above. Under the null hypothesis, the Wald test statistics 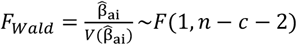 and the exact *P* value of effect (β_ai_) of tested genetic variant can be calculated.

### Permutation test for interaction *cis*-molQTL mapping

Like the standard *cis*-molQTL permutation model (5), we perform direct permutation test for interaction *cis*-molQTL using the following approximate simple linear regression model:

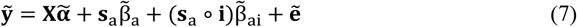

where 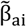 is the interaction effect between the interaction factor and the tested variant corresponding to permuted phenotype values. We focus on the interaction term **s**_a_ ∘ **i** in the permutation test for interaction *cis*-molQTL mapping and define lead variant as the variant with the most significant β_ai_ and/or 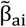. To eliminate the effects of covariates and the effects of **s**_a_, we residualise the interaction term by (**s**_a_ ∘ **i**)_*r*_ = (**s**_a_ ∘ **i**)− 𝕏𝕏^′^(**s**_a_ ∘ **i**)(𝕏^′^𝕏)^−1^ with 𝕏 = (**X, s**_*a*_). The corresponding formula used for residualizing the permuted phenotypes is 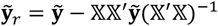, which is computationally intensive and memory intensive when the number of tested variants is large. To avoid the phenotypes residualizing process repeatedly for each variant, we proposed a approximate partial correlation coefficients (APCC) method, that simplify it as 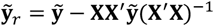 assuming the effect of a single variant on permuted 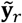 is negligible. To compute the partial correlation, we standardize the 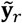 and (**s**_a_ ∘ **i**)_*r*_ to have zero mean and unit sum of squares. Therefore, we simplify the calculation of the partial correlation as 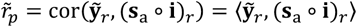

### LMM for dominant molecular QTL mapping

To identify molQTL with dominant effects, we implemented three dominant linear mixed model including LMM(d + A)model corrected with additive GRM (8), LMM(d + D)model corrected with dominant GRM (9), and LMM(d + A + D)and LMM(a + d + A + D)model corrected with both additive and dominant GRMs (10, 11):

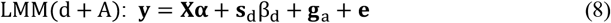

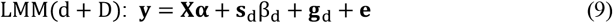

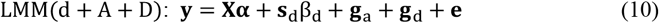

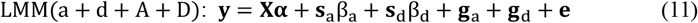

where **s**_d_ is an *n*×1 vector of standardized genotype values in dominant coding (0/1/0 for AA/Aa/aa) for the genetic variant being tested, β_d_ is the variant’s dominant genetic effect, **g**_d_ is an *n*×1 vector of total dominant genetic effects with 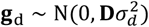 where 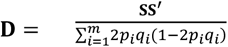 is an dominant GRM, of which **S** is a matrix of standardized genotypes for genome-wide *m* genetic variants and *p*_*i*_ is the MAF at *i*^th^ variant, and the rest of terms are the same as in (1).

To perform association test based on LMM(d + A)or LMM(d + D)models, we can efficiently calculate the 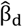 and 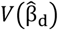 using IDUL the same as formula (3). To perform association test based on LMM(d + A + D)and LMM(a + d + A + D)model, we first estimate the ratio (*θ*_*ad*_) between the additive genetic variance 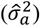 and the dominant genetic variance 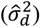 based on

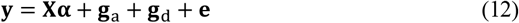

assuming the effect of a single variant on *θ*_*ad*_ is negligible. We obtained the estimates 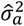 and 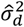 using AI-REML described in below. Given 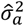 and 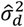, we can write respectively the LMM(d + A + D)model (10) and LMM(a + d + A + D)model (11) as

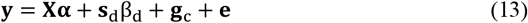

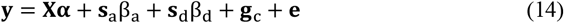

where 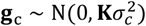 is an *n*×1 vector of total genetic effects combining additive and dominant effects where 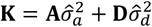 is a combined GRM. To test the β_d_ based on formula (13, 14), we employed the IDUL method as in formula (3) as above. We can obtain estimates 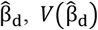, and the corresponding *P*-value from the Wald test statistics 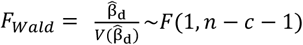. To adjust the multiple testing across molecular phenotypes in dominant *cis*-QTL mapping, we implement permutation test for β_d_ using the same methods as standard LMM(a + A)model (1) and define which phenotypes have significant dominant *cis*-QTL effects.

### Global heritability

#### Additive global heritability

We implement the calculation of the additive global heritability 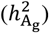 for molecular phenotypes using the following additive genetic model:

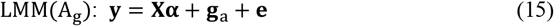

where **y** is an *n*×1 vector of standardized molecular phenotypes, **X** is an *n*×*c* matrix of customized covariates including a column of 1, **α** is the fixed effect corresponding to covariates, **g**_a_ is an *n*×1 vector of total additive genetic effects with 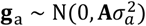 where 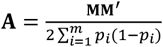 is an additive GRM, of which **M** is a matrix of mean centered genotypes for genome-wide *m* genetic variants and *p*_*i*_ is the MAF at *i*^th^ variant, **e** is *n*×1 vector of residuals with 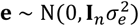 where **I**_*n*_ is an *n*×*n* identity matrix. We estimate the genetic variance 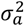 and residual variance 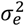 using IDUL algorithm described as follows.

The additive global heritability estimation model assumes **g**_a_ ∼ MVN_*n*_(0, *η τ*^−1^**A**), **e** ∼ MVN_*n*_(0, *τ*^−1^**I**_n_), where *τ*^−1^ is the variance of the residual errors 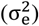, *η* is the ratio between the 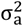 and 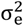, MVN denotes multivariate normal distribution. The eigen decomposition of **A** can be written as **QTQ**^′^ with **QQ**^′^ = **I**_*n*_, where *j*^th^ column of **Q** is an eigenvector, whose corresponding eigenvalue is the *j*^th^ diagonal element of the diagonal matrix **T**. We rotate the formula (15) by multiplying **Q**^′^ to get

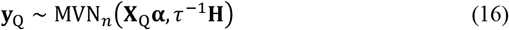

where **y**_Q_ = **Q**^′^**y, X**_Q_ = **Q**^′^**X**, and **H** = *η***T** + **I**_*n*_ . We perform IDUL until *η* converges by repeating:

Step 0: assign a constant to *η*_*i*_ (e.g., 0.1) for the first iteration (*i* = 1) of IDUL.

Step 1: fit formula (3) using weighted least squares with weight **H**^−1^ to obtain the residual **r**.

Step 2: fit **r**^2^ ∼ MVN_*n*_(*µ* + diag(T)γ, *τ*^−1^**H**^2^)using weighted least squares with weight **H**^−2^ to obtain estimates 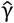 and 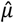.

Step 3: update 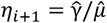 and repeat step 1 to 3 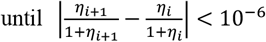 or the number of iterations achieve to the maximum iterations (e.g., 100).

After IDUL converges, we obtain the residual variance estimates 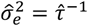, the additive genetic variance 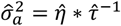, and the 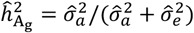.

#### Dominant global heritability

We implement the calculation of the dominant global heritability 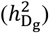 for molecular phenotypes using the following models:

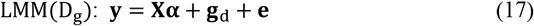

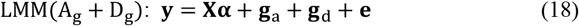

where **g**_d_ is an *n*×1 vector of total dominant genetic effects with 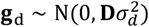 where 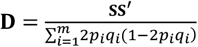 is an dominant GRM. To estimate dominant genetic variance 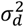 of LMM(D_g_)model (17), we employed IDUL algorithm using method the same as for additive global heritability estimation (i.e., LMM(A_g_)model (15)) and the 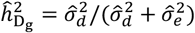. To estimate the 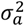 and 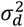 for LMM(A + D)model (18), we employed AI-REML algorithm optimized for molecular phenotypes. For LMM(A_g_ + D_g_)model (18), the variance-covariance matrix of **y** is 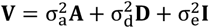. To reduce the repeated calculation across phenotypes for each iteration in AI-REML, we propose a pre-allocated Eigen Decomposition (PAED) method avoiding direct matrix inversion described as follows.

In PAED method, let 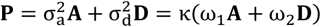 with ω + ω = 1 and κ is a scale factor, we can pre-calculate **P** with different ratios between ω_1_ and ω_2_ (e.g., 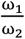 or 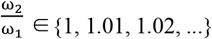). We pre-perform the Eigen Decomposition for all precalcualted **P** above. In the iteration process of AI-REML, we can quickly extract the eigenvalues (**E**_val_) and eigenvectors (**E**_vec_) of **P** with matching α and β for calculating 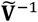. We can easily to calculate 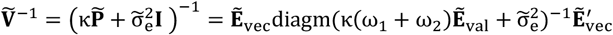 with 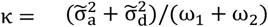. Finally, we obtain the 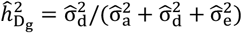.

### *Cis* heritability

#### Additive cis heritability

For estimating the contribution of additive genetic effects 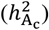 for molecular phenotypes, we implement three genetic models described as follows:

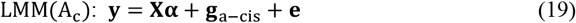

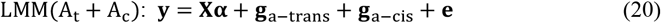

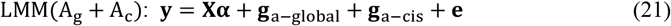

where 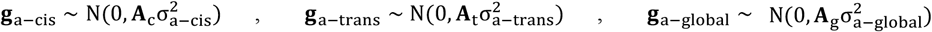 where **A**_c_, **A**_t_, and **A**_g_ are additive *cis*-GRM, *trans*-GRM, and global-GRM constructed using genotypes from *cis*-region variants, *trans*-region variants, and genome-wide variants, respectively. By default, we define the *cis*-region as the ± 1Mb region surrounding the transcription start sites (TSS) of the target molecular phenotype, whereas the *trans*-region refers to chromosomes distinct from the one where the target molecular phenotype is situated.

For LMM(A_c_)model (19), we estimate the residual variance 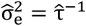, the *cis* additive genetic variance 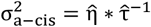 using IDUL the same as for the LMM(A_g_)model (15) and obtain the 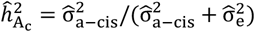.

For LMM(A_t_ + A_c_)model (18), we proposed a *cis* low-rank approximation (*cis*LRA) method to implement AI-REML optimized for molecular phenotypes. In *cis*LRA, we assume that the *cis*-GRM can be represented with low-rank matrices due to high linkage disequilibrium in *cis*-region. We decomposed *cis*-GRM into low-rank matrices multiplication using Partial Singular Value Decomposition (PSVD) with **A**_c_ ≈ **UΣΛ**^′^, where **Σ** is a *k*×*k* diagonal matrix, **U** and **Λ** are *n*×*k* matrices (*k* ≪ n). Based on the Woodbury Matrix Identity (**A** + **UΣΛ**^′^)^−1^ = **A**^−1^ − **A**^−1^**U**(**Σ**^−1^ + **Λ**^′^**A**^−1^**U**)^−1^**Λ**^′^**A**^−1^, we danote 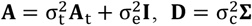, where **A**^−1^ can be quickly calculated using Eigen Decomposition. We calculate the variance-covariance matrix of **y** in the iteration process of the AI-REML as 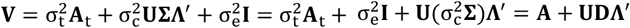. The inversion of **V** is 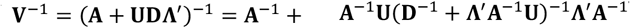, where (**D**^−1^ + **Λ**^′^**A**^−1^**U**)is a *k*×*k* low-rank matrix that can be inversed quickly. We obtain the 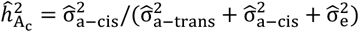.

The method for estimating variance components of the LMM(A_g_ + A_c_)model (21) is the same as for the LMM(A_t_ + A_c_)model (20) and the 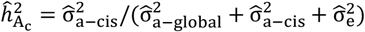.

#### Dominant cis heritability

For estimating the contribution of dominant genetic effects 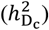 for molecular phenotypes, we implement three genetic models described as follows:

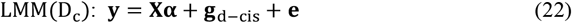

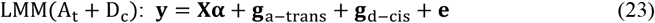

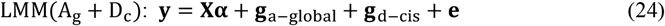

where 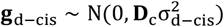 where **D**_c_ is dominant *cis*-GRM constructed using genotypes from *cis*-region variants. We estimate the variance components for LMM(D_c_)model (22), LMM(A_t_ + D_c_)model (23), and LMM(A_g_ + D_c_)(24) using the same methods as for LMM(A_c_)model (19), LMM(A_t_ + A_c_)model (20), and LMM(A_g_ + A_c_)(21), respectively. We obtain the 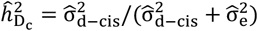, the 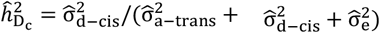, and the 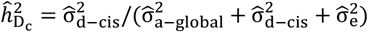 for LMM(D_c_)model (22), LMM(A_t_ + D_c_)model (23), and LMM(A_g_ + D_c_)(24), respectively.

### *Trans* heritability

We implement the calculation of the additive *trans* heritability 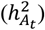 and dominant *trans* heritability 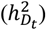 for molecular phenotypes using the following models:

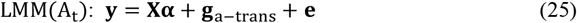

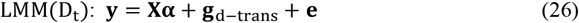

where 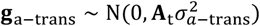 and 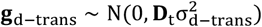 where **A**_t_ and **D**_c_ are the additive *trans*-GRM and dominant *trans*-GRM constructed using genotypes from *trans*-region variants of target molecular phenotype. We estimate the variance components of LMM(A_t_)model (25) and LMM(D_t_)model (26) using IDUL the same as for LMM(A_g_)model (13). We obtain the 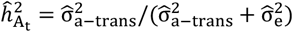 and 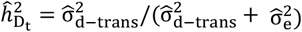 for LMM(A_t_)model (25) and LMM(D_t_)model (26), respectively.

### Summary of real data

We collected three public datasets including GIAD042^27^, MAGE^25^, and GEUVADIS^26^ described as below.

GIAD042 comprises 300 pigs from three distinct subpopulations: Duroc (n=100), Yorkshire (n=100), and Landrace (n=100). The Duroc (DUR) individuals are offspring from 33 sows and 10 boars, Yorkshire (YOR) individuals from 84 sows and 43 boars, and Landrace (LAN) individuals from 74 sows and 18 boars. The genotype data of all 300 individuals were obtained from the GigaScience GigaDB database (http://dx.doi.org/10.5524/102388). Following quality control across all 300 pigs, we retained 15,495,927 biallelic variants with MAF≥5% and MCA≥6 in 18 autosomes. Additionally, we obtained 900 RNA-Seq samples from three tissues (duodenum, liver, and muscle). To exclude lowly expressed genes (i.e., TPM < 0.1 and/or raw read counts < 6 in more than 80% of samples within each of tissues), we retained a total of 14,007, 12,512, and 9,164 expressed autosomal genes in duodenum, liver, and muscle, respectively. We normalized the raw counts by trimmed mean of M values (TMM) within each tissue and used the inverse normal transformed TMM for subsequent analyses.

MAGE (Multi-ancestry Analysis of Gene Expression) comprises 731 unrelated individuals from five distinct human populations including AFR (n=196), AMR (n=113), EAS (n=141), EUR (n=142), and SAS (n=139). We obtained the genotype data of all 731 individuals from the 1000 Genomes Project (1KGP) samples (http://ftp.1000genomes.ebi.ac.uk/vol1/ftp/data_collections/1000G_2504_high_coverage/working/20201028_3202_phased/). After quality control across all 300 individuals, we kept 14,267,797 biallelic variants with MAF≥1% and MAC≥6 in 22 autosomes. We obtained 731 RNA-Seq samples from EBV transformed lymphoblastoid cells lines (LCLs). To eliminate the lowly expressed genes using the same rules as did in GIAD042, we obtained a total of 19,539 expressed autosomal genes and used their inverse normal transformed TMM for subsequent analyses.

GEUVADIS (Genetic European Variation in Disease) comprises 462 unrelated individuals from five distinct populations: CEU, FIN, GBR, TSI and YRI. The genotype data were sourced from the 1000 Genomes Project. After quality control across all 462 individuals, we retained 10,313,747 biallelic variants with MAF≥1% and MAC≥6 in 22 autosomes. We obtained 462 RNA-Seq samples from LCLs. To eliminate the lowly expressed genes using the same rules as did in MAGE, we obtained a total of 17,703 expressed autosomal genes and used their inverse normal transformed TMM for subsequent analyses.

### Genome-wide linkage disequilibrium

To evaluate the pattern of genome-wide linkage disequilibrium, we split the genotype data into different sub-populations of GIAD042 and MEGA datasets. For each of all sub-populations, we kept SNPs with MAF>5% for subsequent analysis. We computed the intra- and inter-chromosomal mean LD using GEAR^50^.

### Genotype data used in simulations

To assess the performance of OmiGA in comparison to other methods, we conducted a comprehensive simulation study, utilizing both populations consisting of related individuals and populations comprising only unrelated individuals. We used the WGS data from two public populations including the GIAD042^27^ and MEGA^25^ datasets above mentioned, where the GIAD042 is a pig population with close relatedness and the MAGE is a human population without related individuals. We used PLINK v2.0^53^ and BCFtools^54^ to carry out quality control on the datasets. For both datasets, we retained biallelic autosomal variants with call rate ≥95%, minor allele frequency (MAF) ≥5%, and minor allele count (MAC) ≥6. We phased and imputed the missing genotypes using Beagle5.2^40^ without reference panel. We conducted LD pruning using the flag --indep-pairwise 50 5 0.25 in PLINK v2.0^53^, resulting in 770,243 and 492,797 variants in the GIAD042 and the MAGE, respectively, for the downstream simulations. Given the significant difference of the degree of kinship among individuals, we take the simulations using the GIAD042 and the MAGE named as SIM-CLOSE and SIM-UNREL below, respectively.

### Schemes for simulating gene expression

To evaluate the performance of OmiGA in *cis*-eQTL mapping, we simulated genes with causal variants within *cis*-region in SIM-CLOSE and SIM-UNREL. We defined 1Mb upstream and downstream of the transcription start site (TSS) of each gene as the *cis*-region. Based on the genome coordinate of genes from Ensembl, we used only those genes located on chromosomes 1 and 2 and have at least 30 *cis*-variants for the simulations.

#### *cis*-eQTL simulation

Various study designs suggest that approximately 60% to 90% of the genetic variance in expression can be attributed to *trans*-acting variation^55–58^. However, *trans*-eQTLs are notoriously difficult to find in humans is mainly due to effect sizes are considerably small, making it more difficult to achieve statistical significance compared to *cis*-eQTLs^59^. Fisher’s polygenic model^60^ assumes that a trait is controlled by numerous loci, each of which exerting a weak effect. The framework can be utilized to simulate the causal effects within *trans*-regions. We can denoted the contributions from *cis* (local) and *trans* (distant) regions to the gene expression (***y***) of an individual as:

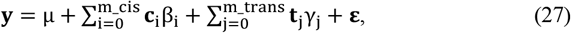

where μ is the overall mean, **c** and **t** represent the genotypes for causal variants in the *cis* and *trans*-regions, respectively, β_i_ and γ_j_ represent their corresponding effect size, m_cis and m_trans represent the number of causal *cis*-variants and *trans*-variants, respectively, and **ε** is the residual error. In the *trans*-region, the cumulative effects of these causal variants can be approximated as a normally distributed random variable (polygenic_*trans*_) in accordance with the Central Limit Theorem^61^.

Therefore, we simplify the formula (27) to simulate the expression levels for each gene as follows:

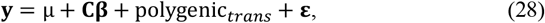

where **y** ∈ ℝ^*n*^ is a vector representing the simulated gene expression levels across *n* individuals, **C** is the matrix the standardized genotypes of *m*_*cis* causal variants in the *cis*-region of target gene, **c**_i_ = s_i_ − 2p_i_, where **s**_i_ ∈ {0,1,2} and p_i_ represents the MAF of i^th^ causal variants, the vector **β** represents the effect sizes of casual variants on gene expression, the polygenic_*trans*_ obey the multivariate normal distribution of *N*(**0**, σ^2^ **K**^**⋆**^)with **K**^**⋆**^ denotes the kinship matrix constructed using the LD-independent variants in *trans*-region and the *trans* genetic variance 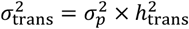, the residual error 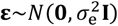 with 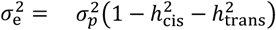, where the 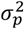 is the phenotypic variance, 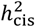 is the *cis*-heritability, and 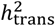 t*rans*-heritability.

Many evidences suggest that the genetic architecture of *cis*-regions is sparse for many genes^62,63^. To simulate genes with causal variants in *cis*-region, we sample the number of causal variants on the basis of a prior probability table (Supplementary Table 1) identical to that in ^24^, which summarizes findings from GTEx’s independent *cis*-eQTL analysis^3,42^. For each of casual variant in *cis*-region, we sample its effect β_i_ from a normal distribution 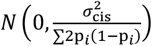, where 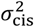 is the *cis* genetic variance and p_i_ represents the MAF of i^th^ causal variants. Let 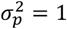 for each gene, we obtained the *cis* genetic variance 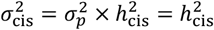. We then scaled the sampled effect **β**_i_ of i^th^ casual variant by multiplying the factor 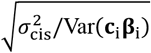. To simulate genes with various *cis*-heritability, we generate the *cis*-heritability from a scaled Gamma distribution of 0.1 * rand(*Γ*(0.6,2))and constraint it between 0.005 and 0.6, where rand(*Γ*(0.6,2))represents a random number sampled from the Gamma distribution with a shape parameter *α* = 0.6 and a scale parameter *β* = 2.

To mimic the part of polygenic_*trans*_, we further use two constraints including that (1) the proportion of total genetic variance explained by *cis*-region less than 90%, (i.e., 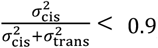), and (2) the sum of *cis*- and *trans*-heritability at the gene level should less than 0.8 (i.e., 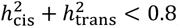). Given a *cis*-heritability, we can obtain the minimum and maximum of *trans*-heritability, denoted as 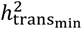 and 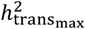, respectively. We calculate the *trans*-heritability using the formula: 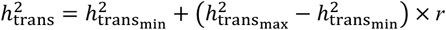 with 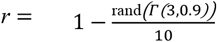, where rand(*Γ*(3,0.9))represents a random number sampled from the Gamma distribution with a shape parameter *α* = 3 and a scale parameter *β* = 0.9 and we restrain the *r* ranged from 0 to 1. Finally, we obtained the polygenic_*trans*_ from the multivariate normal distribution of 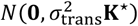 and scale it by multiplying the factor 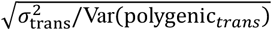.

Based on the strategies above, we simulated 3,556 eGenes and 1,879 non-eGenes in the SIM-CLOSE, and 2,171 eGenes and 1,144 non-eGenes in the SIM-UNREL, where the eGenes and the non-eGenes are the genes simulated with at least one causal *cis*-variant and zero causal *cis*-variant, respectively. All the simulations were repeated 50 times.

#### Interaction *cis*-eQTL (*cis*-ieQTL) simulation

To evaluate the performance of OmiGA in context-dependent *cis*-eQTL (i.e., *cis*-ieQTL) mapping, we simulated genes with ancestry-specific effects for causal variants within *cis*-region in SIM-CLOSE and SIM-UNREL. We first estimated the proportion of the Yorkshire ancestry of 300 individuals from GIAD042 populations and the AFR ancestry of 731 individuals from MAGE populations using ADMIXTURE v1.3.0 based on the genotype data.

To simulate genes with ancestry-specific effects for causal variants, we used the following formula to generate the expression levels (**y**) for each gene:

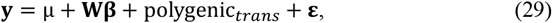

**y** = μ + **Wβ** + polygenic_*trans*_ + **ε**, (29) where μ, polygenic_*trans*_, and **ε** is the same as those in formula (28), **W** is the matrix of interaction term between genotype and the proportion of ancestry with **W** = **C** ∘ **a**, of which **C** is the matrix the standardized genotypes of *m*_*cis* causal variants in the *cis*-region of target gene, **c**_i_ = s_i_ − 2p_i_, where **s**_i_ ∈ {0,1,2} and p_i_ represents the MAF of i^th^ causal variants and **a** ∈ ℝ^*p*^ is the vector of the proportion of ancestry across individuals, **β** is the sampled effect sizes of causal variants follows the normal distribution of *N*(0,1). We further scale the sampled effect sizes **β** by multiplying the factor 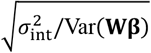, where 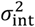 is the genetic variance explained by the interaction between genotype and ancestry. We sample the number of causal variants on the basis of a prior probability table (Supplementary Table 2) and restrain the proportion of phenotypic variance explained by interaction term 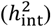 to a range of 0.005 to 0.3. All other settings are consistent with formula (28), wherein 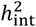 substitute for 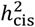.

Based on the strategies above, we simulated 2,459 ieGenes and 2,976 non-ieGenes in the SIM-CLOSE, and 1,513 ieGenes and 1,802 non-ieGenes in the SIM-UNREL, where the ieGenes and the non-ieGenes are the genes simulated with at least one causal *cis*-variant and zero causal *cis*-variant, respectively. All the simulations were repeated 50 times.

### Performance comparison for molQTL mapping

To quantify the performance of different methods, we computed several metrics including the genomic inflation factor (λ), false discovery rate (FDR), F1-score, and true positive rate (TPR). Median λ is defined as the median chi-squared statistic divided by its expected value at the null variants. The FDR is defined as the proportion of simulated non-eGenes within these genes that were reported as significant (for example, with a *P*-value < 0.05). The F1-score is computed by the formula 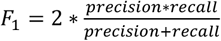, where 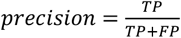 and 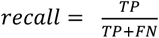, in which TP, FP, and FN represent the number of true positive, false positive, and false positive genes, respectively. The TPR is defined as the proportion of simulated eGenes that can be discovered under 5% FDR.

### Statistics & Reproducibility

No statistical method was used to predetermine the sample size. The details of data exclusions for each specific analysis are available in the Methods section. For all the boxplots, the horizontal lines inside the boxes show the medians. Box bounds show the lower quartile (Q1, the 25^th^ percentile) and the upper quartile (Q3, the 75^th^ percentile). Whiskers are minima (Q1 − 1.5 × IQR) and maxima (Q3 + 1.5 × IQR), where IQR is the interquartile range (Q3-Q1). Outliers are shown in the boxplots unless otherwise stated. The experiments were not randomized, as all the datasets are publicly available and from observational studies. The Investigators were not blinded to allocation during experiments and outcome assessment, as the data are not from controlled randomized studies.

## Data availability

The individual-level genotype and gene expression data of MAGE are publicly available from http://ftp.1000genomes.ebi.ac.uk/vol1/ftp/data_collections/1000G_2504_high_coverage/working/20201028_3202_phased/ and https://doi.org/10.5281/zenodo.10072081, respectively. The individual-level genotype and gene expression data of GEUVADIS are publicly available from http://ftp.1000genomes.ebi.ac.uk/vol1/ftp/release/20130502/ and https://www.internationalgenome.org/data-portal/data-collection/geuvadis. The genotype and RNA-Seq data of GIAD042 are obtained from http://dx.doi.org/10.5524/102388 and the SRA Accession: PRJEB58031. The genotype and gene expression data of 34 pig tissues from the pilot phase of PigGTEx project are available from the PigGTEx portal (https://piggtex.farmgtex.org/). All the full summary statistics of molQTL mapping generated from this study are available at our portal (https://omiga.bio/).

## Code availability

OmiGA is available at https://omiga.bio/.

## Notes

https://omiga.bio

### Competing Interest Statement

The authors have declared no competing interest.

